# Unifying error and reward action learning: a cerebello-basal ganglia theory

**DOI:** 10.64898/2026.08.22.745751

**Authors:** Michele Garibbo, Carolina Filipe, Laurence Aitchison, Rui Ponte Costa

## Abstract

Learning depends on both reward- and error-based feedback, yet how the brain integrates these distinct signals to guide behaviour remains fundamentally unclear. Here, using a normative computational framework, we derive credit assignment rules for both reward-based learning (RBL) and error-based learning (EBL). In contrast to existing dual-policy accounts, our approach demonstrates that RBL and EBL updates can be reformulated into a shared action-gradient space that directly updates a single downstream policy. First, we map this action-gradient framework onto a systems-level account of coordinated interactions between the basal ganglia, cerebellum, and cortex. The model reproduces key behavioral features across both learning regimes, generates experimentally testable predictions, and provides a unified computational account of motor deficits observed in patients with cerebellar and basal ganglia disorders. Together, our work offers a normative, brain-wide framework for how distributed brain systems integrate reinforcement and error-driven feedback toward a common behavioral objective.

## Introduction

Motor commands typically generate two forms of feedback: sensory outcomes, reflected in the activity of sensory systems such as vision and proprioception, and rewards, which represent subjective evaluations of the utility or success of those outcomes. Historically, these feedback signals have been investigated within largely distinct learning frameworks: error-based learning (EBL) and reward-based learning (RBL) ^1–3^.

RBL focuses on learning from rewards and has been linked to dopamine in the basal ganglia, aligning with actor-critic architectures in reinforcement learning (RL) theory ^4–8^. In contrast, EBL accounts for learning from sensory feedback and has been associated with the cerebellum, consistent with principles of supervised learning ^9–11^.

Crucially, RBL and EBL have traditionally been viewed as distinct and largely independent learning systems ^2,3,9,12^. This perspective aligns with the long-standing anatomical view that the cerebellum (CB) and the basal ganglia (BG) form separate, parallel channels through the thalamus ^13–15^. This separation has led to the implicit assumption that reward and error feedback drive the learning of separate motor policies — one mediated by the basal ganglia and another by the cerebellum ^2,8,16,17^. However, this view raises several unresolved questions. Most notably, it is unclear how the brain coordinates learning across two independent policies to solve the same task. For instance, it is not obvious how information acquired through rewards transfers to an error-driven policy, or vice versa — nor how the brain might keep the two policies synchronized over time (we investigate this in Supplementary material). Furthermore, it is not evident why maintaining and updating two separate policies would be computationally advantageous, especially given robust evidence that the brain can integrate reward and sensory feedback during learning within a single task context ^1,2,18,19^.

Recent anatomical and functional studies further challenge the notion of strict segregation between the cerebellum and basal ganglia. A growing body of work demonstrates tight interactions between these structures ^14–16,20–25^. In particular, disynaptic projections from the cerebellum to the dorsal striatum (i.e., caudate and putamen) via the intralaminar nuclei of the thalamus (ILN) have been well documented ^14,20,21,24^. Moreover, cerebellar activity has been shown to influence striatal function and plasticity, with clear behavioral consequences ^21,24–26^, suggesting a more integrated role for the cerebellum in modulating learning within the basal ganglia.

Motivated by these findings and building on action gradient theory ^27^, we propose a novel learning framework that explicitly integrates sensory and reward error signals, reflecting the functional interplay between cerebellar and basal ganglia circuits. By expressing sensory errors and reward prediction errors within a common action-gradient space, the framework enables their principled combination — a goal that is difficult to achieve when these signals are treated as independent teaching signals. Their weighted integration yields a unified error signal that can drive plasticity in downstream motor circuits responsible for generating behavior. To support this account, we propose a system-level implementation consistent with accumulating evidence for functional interactions between the cerebellum and basal ganglia during learning ^15,21–24,26^. We demonstrate that this framework accounts for a broad range of behavioral phenomena associated with reward-based learning (RBL), error-based learning (EBL), and their interaction ^2,28–30^, including characteristic impairments observed following cerebellar or basal ganglia dysfunction.

We next introduce a cerebellar network that computes the EBL-derived component of the action gradient. Although this role is consistent with the widely held view that the cerebellum learns internal models of the world, it departs from the classical assumption that the cerebellum primarily implements a forward model. We show that the proposed network captures several hallmark features of motor adaptation, including savings and interference, while also providing a mechanistic account of classical cerebellar deficits such as motor ataxia.

Crucially, the framework integrates this cerebellar contribution to EBL with the established role of dopaminergic reward prediction errors in RBL ^4–6,8^, thereby unifying two learning processes that have traditionally been studied in isolation. We conclude by deriving several experimentally testable predictions concerning: (1) a potential role for the cerebellum in shaping synaptic plasticity within the dorsal striatum e.g., ^24^; (2) the contribution of reward to motor memory consolidation e.g., ^18,31–34^; and (3) the interaction between RBL and EBL during the acquisition of novel motor skills.

## Results

Here, we first introduce the action-gradient learning framework and outline its implementation within known brain systems. We then demonstrate how this framework accounts for a diverse range of experimental findings in motor learning, as well as region-specific deficits in motor function.

### Action-gradient framework for reward- and error-based learning

Our framework requires the formulation of two complementary learning rules corresponding to reward-based learning (RBL) and error-based learning (EBL; Fig. 1). To express these updates within a common mathematical framework, we adopt the terminology of reinforcement learning (RL). We model motor commands as actions, *a*_*t*_, generated by a policy *π*_*ϕ*_ parameterised by the activity of a motor circuit with parameters *ϕ*. The policy maps latent motor representations, *h*_*t*_, onto actions according to 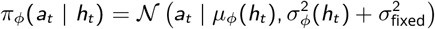. Following previous work ^6,27^, we assume that actions are sampled from a Gaussian distribution whose mean, *µ*_*ϕ*_(*h*_*t*_), and standard deviation, *σ*_*ϕ*_(*h*_*t*_), are determined by the policy parameters *ϕ*. The additive term *σ*_fixed_ introduces an irreducible component of motor variability, capturing the persistent background noise observed in human motor behaviour ^29^ (see Methods for details).

**Figure 1.**
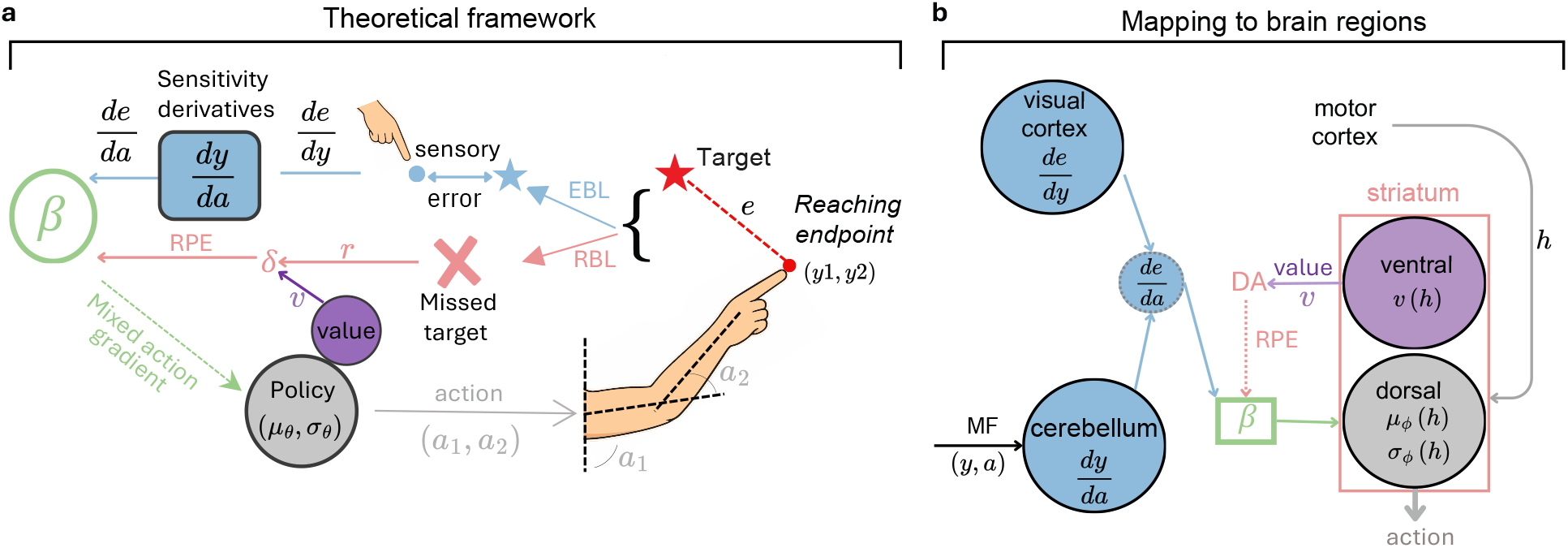
Action-gradient theory: conceptual schematic and mapping to systems neuroscience. **a**, Schematic of the proposed action-gradient learning framework for endpoint arm reaches. The policy (grey circle) outputs an action that determines the arm endpoint location. From the resulting endpoint, two forms of feedback are computed e.g., ^2^: (1) a sensory error given by the distance between the arm endpoint and the target location, and (2) a reward indicating whether the reach is on or off target. The sensory error drives error-based learning (EBL; blue pathway), which requires estimates of the sensitivity derivatives, 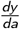, to determine how the action should be modified to reduce the sensory error. The reward feedback generates a reward prediction error (RPE), driving reward-based learning (RBL; red pathway). Crucially, both forms of feedback can be expressed as action gradients (see main text) and combined through a *β*-weighted sum, yielding a mixed learning signal that drives policy learning. **b**, Systems-level implementation of the proposed action-gradient learning framework within cerebellar-basal ganglia-cortical circuitry ^25^. The dorsal and ventral striatum encode the policy and critic, respectively e.g., ^4,5,8^. The motor cortex provides latent motor representations, *h*_*t*−1_, to the striatum, supporting rich action representations. The cerebellum provides estimates of the sensitivity derivatives, 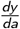, which, together with afferent sensory errors, are used to compute the EBL action gradient, 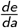. Dopaminergic projections from midbrain dopamine neurons in the ventral tegmental area (VTA) convey reward prediction errors, δ_*t*_, to the dorsal striatum, providing the signals required to compute the RBL action gradient. Thus, EBL- and RBL-derived action gradients converge in the dorsal striatum, enabling policy learning driven jointly by sensory and reward feedback. RPE: reward prediction error. VTA, ventral tegmental area.

Within this framework, RBL and EBL specify how the policy parameters *ϕ* are updated in response to reward signals, *r*_*t*_, and sensory error signals, *e*_*t*_, respectively. Through repeated updates across trials, these learning processes adjust the policy to maximise reward and minimise sensory error, thereby improving motor performance (see Methods for detailed derivations of the RBL and EBL update rules). Throughout the main text, we present parameter updates, Δ*ϕ*_*t*_, for the simple case of one-dimensional actions within a one-dimensional sensory space. This notation is adopted solely for clarity of exposition; the framework extends naturally to higher-dimensional action and sensory spaces, as illustrated in several of the motor-learning tasks considered below.

The RBL update (see Methods) can be factorised into two terms, 1) an ‘action gradient’ term, 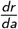, which defines a learning signal, instructing how, given reward, the current action should change to improve performance and, 2) a parameter update term, 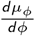, which drives (synaptic) plasticity in a downstream action-encoding motor area ^6,27^. Therefore, RBL is given by

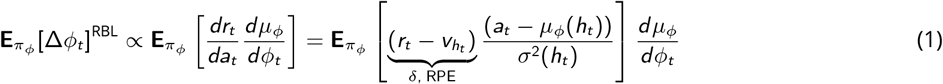

where 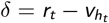 denotes the classical reward prediction error (RPE), while *σ* denotes the policy estimated variance (see Methods for details and derivation). It is important to stress that the reward signal *r* can be discrete (e.g., encoding success/failure signal) or continuous (e.g., encoding how far we are from a desired goal), depending on the task. Crucially, the standard EBL update ^27,35^ can also be factorised into two equivalent terms, 1) the teaching signal (‘action gradient’) term given by, 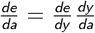, and 2) a parameter update term given by, 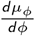. Here, the sensory error *e* is typically computed as the (squared) distance between a target, *y*^∗^ and the actual movement outcome, *y* (i.e., the task error, *e* = (*y*^∗^ − *y*)^2^). Since 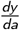 determines how small changes in *a* influence the movement outcome *y*, we refer to these as *sensitivity derivatives*.

This equivalence between the RBL and EBL synaptic weight updates allows us to perform a weighted combination of the RBL and EBL learning signals in the same action gradient space to jointly drive plasticity of downstream action-encoding motor areas. This ‘unified’ update can be expressed as follows (we removed the expectation over the policy *π*_*ϕ*_ for simplicity),

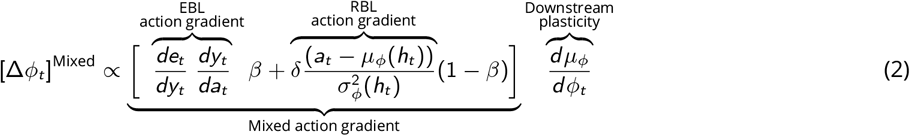

where Δ*ϕ*_*t*_ denotes the change to the policy parameters (e.g., synaptic plasticity at an action-encoding motor area) at trial *t* as a function of both RBL and EBL learning signals. *β* ∈ [0, 1] models the trade-off between the RBL and EBL action gradient terms (i.e., learning signals). By controlling this trade-off, *β* should depend on the quality of sensory information. This role of *β* is in line with previous work showing that the magnitude of EBL-driven adaptation negatively correlates with sensory feedback uncertainty ^2,30,36,37^. When no sensory feedback is available and only a reward signal is provided, the EBL action gradient cannot be computed. In this scenario, learning must rely entirely on the RBL action gradient (i.e., *β* = 0) ^38,39^. At the opposite extreme, when ‘perfect’ sensory information is available, the EBL action gradient offers the most effective learning strategy (i.e., *β* = 1) ^27,40,41^. Note that we also optimize the policy parameters based on its variance 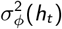, which results in similar updates (see Methods).

Beyond these extremes, our action-gradient model (Eq. 2) predicts that the brain smoothly integrates reward-based (RBL) and sensory feedback into a single, unified teaching signal to drive plasticity in downstream motor areas encoding the policy, *ϕ*. This contrasts with traditional accounts that treat RBL and EBL (error-based learning) as separate processes operating on distinct policies ^2,16^. Maintaining separate policies seems unlikely, as modifying one (e.g., when only reward is available) would require complex, non-trivial adjustments to the other to ensure coherent interaction (see Supplementary Material). Instead, by merging RBL and EBL into a unified signal, our model parsimoniously explains a broad range of phenomena, including motor deficits in cerebellar and basal ganglia dysfunctions, long-term memory consolidation via RBL, and RBL-EBL interactions during de novo motor learning.

Furthermore, in the Supplementary Material (Fig. S3), we show that our learning framework is more plausible than the traditional view of separate RBL and EBL policies, which requires continuous synchronization as sensory and reward feedback change.

### Mapping framework with cerebello-cortico-basal ganglia systems

In the classical actor-critic model of basal ganglia learning, the dorsal and ventral striatum function as the policy (actor) and the critic, respectively ^4,5,8^. Both regions receive midbrain dopaminergic projections that convey reward prediction errors (RPEs), denoted as δ ^4,42^. Within this framework, the ventral striatum uses these RPEs to update the expected value of states or actions. Meanwhile, the dorsal striatum uses the same RPE signal to drive cortico-striatal plasticity according to an RBL action gradient, ultimately learning to select optimal actions ^4,6,43^. Crucially, the inputs to these dorsal circuits are provided by cortico-striatal synapses, which convey latent action representations *h* (such as motor plans) from the primary motor cortex (M1) to the striatum ^9,44,45^.

Our action gradient theory proposes to complement this classical view of dopamine-driven learning in the basal ganglia, with error-driven learning, thereby providing a principled and integrated explanation for the growing evidence of cerebellar (CB) driven signals in the basal ganglia (BG) ^15,21–26^. Specifically, our model (Eq. 2) predicts that dorsal striatal plasticity (i.e., the policy weights, *ϕ*) is not only driven by dopamine, but also by cerebellar teaching signals, reflecting a combination of dopamine-driven RBL and cerebellar-driven EBL feedback (Fig. 1).

For EBL to take place, two quantities must be computed, 1) the sensitivity derivatives, 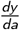, and 2) the ‘directed’ sensory error 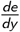 (Eq. 9). We propose that the cerebellum learns to compute the sensitivity derivatives 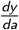 ^46^, by tracking how (small) changes to actions, Δ*a*, impact the resulting sensory outcomes, Δ*y* (see Methods for more details). We think that the cerebellum is well placed to learn the sensitivity derivatives, 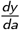, given its established role in encoding internal models of action-sensory interactions ^10,12,47–50^, which are critical to EBL ^10,51–53^. Conversely, we assume that the (sensory) cortical areas estimate the ‘directed’ sensory error component, 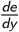, depending on the sensory modality in which the sensory outcomes of the task are perceived (e.g., visual, tactile, etc.). For instance, photoinhibition of the somatosensory cortex seems to impair EBL of (novel) force field perturbations, while leaving previously learned motor commands unaffected ^54^.

Finally, the cerebellum and the cortical predictions converge to the dorsal striatum, the former via targeted disynaptic projections going through the intralaminar nuclei of the thalamus (ILN) e.g., ^21,25,26^, while the latter via the well-known (sensory) cortical projections to the dorsal striatum ^55^. Together, the two signals provide the necessary information to compute the EBL action gradient, driving dorsal striatal plasticity accordingly. This is consistent with the evidence that cerebellar drives dorsal striatal plasticity together with cortical projections ^24^.

In order to combine both EBL and RBL, our action-gradient theory uses a *β*-weighting Eq. 2. We speculate this mechanism could take place implicitly, through (synaptic) competition mechanisms driven by the magnitude of each gradient or, explicitly, with the cerebellum directly controlling midbrain dopamine release at the striatum, as some evidence indicates ^56–59^.

In summary, the proposed circuit highlights how learning in the dorsal striatum (policy) may be driven by a mixture of cerebellar- and dopamine-driven signals, reflecting unified RBL and EBL, as described by the action-gradient model in eq. (2). In support of this proposal, cerebellar projections appear to target medial spiny neurons (dorsal striatal), which also possess D1 and D2 dopamine receptors ^21^, thus allowing cerebellar and dopamine feedback to converge within the same neurons.

### Action-gradient framework predictions of behavioural data

Here, we demonstrate that our action-gradient framework accounts for motor adaptation during arm-reaching tasks that integrate reward-based (RBL) and sensory error-based (EBL) feedback. Following the paradigm of Izawa and Shadmehr ^2^ (Fig. 2a), the model learns to reach a target under a visuomotor perturbation that gradually increases over the trials up to 8°. We evaluate the model under three feedback conditions:

**Figure 2.**
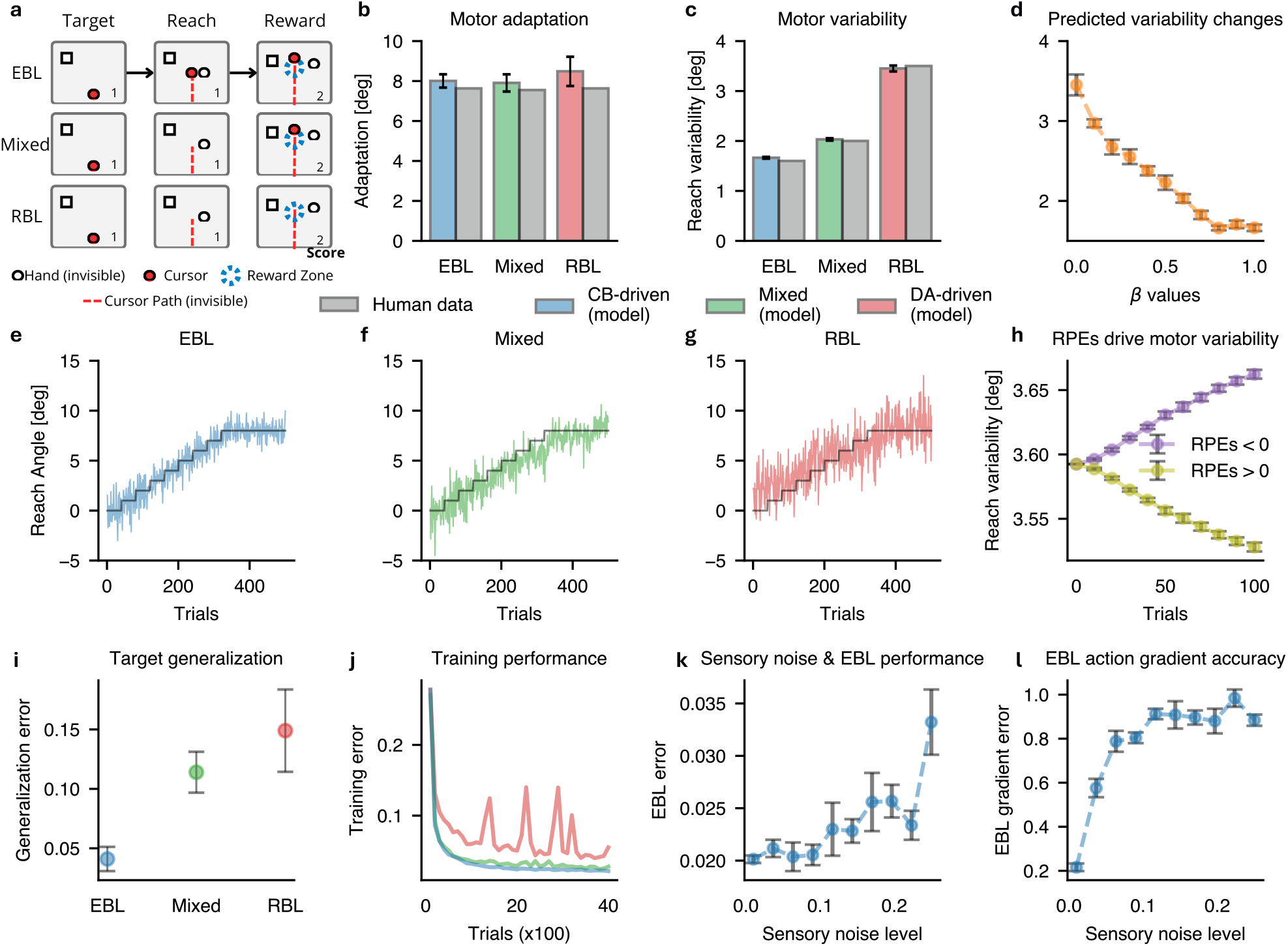
RBL-EBL action gradient framework accounts for human motor learning observations. **a**, Task diagram for the conditions studied experimentally, figure adapted from Izawa and Shadmehr ^2^. For EBL, feedback includes the cursor trajectory, the final cursor endpoint, and the reward outcome. For the mixed condition, feedback is limited to the final cursor endpoint and the reward outcome, with no trajectory information (i.e., reduced visual feedback). For RBL, feedback is limited to the reward outcome alone (i.e., whether the cursor reached the reward zone). **b**, Motor adaptation comparison between our action-gradient model and human experimental data during EBL (blue), RBL (red) and a mixture of the two (green). Human data were re-plotted from Izawa and Shadmehr ^2^. CB: cerebellum, DA: dopamine **c**, Similar to (c) but for motor variability. **d**, Model predicted change in motor variability as a funcion of *β*. **e-g**, Reaching angle adaptation across EBL (e), Mixed (f) and RBL (g), the black line represents the required change in reaching angle. **h**, Model predicted change in motor variability during RBL in response to either negative (purple) or positive (yellow) reward prediction errors. **i,j**, Action-gradient model’s generalisation performance (i) and training errors (j) across EBL, RBL and a mixture of the two. **k,l**, EBL performance and EBL action gradient accuracy (l) across different levels of sensory noise. Error bars are calculated across five random seeds.

1. **EBL condition** (ERR in the original study): Full visual trajectory feedback is provided alongside terminal binary reward (success/failure).
2. **RBL condition** (RWD in the original study): Only terminal binary reward is provided, withholding visual information to prevent error-based updating.
3. **Mixed condition** (EPE): Binary reward is paired with restricted visual feedback, heightening sensory uncertainty so that both reward and visual cues contribute to learning.

Below, we show that our model accounts for human motor adaptation, variability, and generalization across all three conditions. This is achieved by tuning *β* in Eq. 2, which regulates the relative contributions of the RBL and EBL action gradients.

The reward-only RBL condition is captured by setting *β* = 0, where plasticity is driven exclusively by dopamine (DA-driven) without cerebellar involvement due to the absence of visual feedback. Conversely, the EBL condition corresponds to *β* = 1, where plasticity is mediated entirely by sensory errors and the cerebellum.

This indicates that when precise sensory information is available (i.e., low sensory uncertainty), human motor adaptation relies predominantly on cerebellar-driven EBL, consistent with the fact that an accurate EBL action gradient provides a richer teaching signal than RBL e.g., ^27,40,41^. Finally, setting *β* = 0.4 yields the best fit for the Mixed condition (EPE), where both rewards and sensory errors drive learning, suggesting that humans dynamically weight both action gradients when faced with sensory uncertainty. We report these results in detail below.

#### Motor adaptation and motor variability

Our action-gradient model predicts human motor variability across the RBL, EBL, and Mixed conditions (Fig. 2b) while also capturing the amount of motor adaptation in each condition (Fig. 2a). Note that, although the model predicts full adaptation to the 8° perturbation in all three conditions, humans slightly under-adapt throughout, possibly because of a small bias toward one side of the target e.g., ^35^. In the RL literature, the RBL action gradient is known to have a higher variance (i.e., noisier) than the equivalent EBL gradient ^27,40^. Our model therefore suggests that changes in human motor variability across conditions may reflect the variability of the underlying action-gradient estimates (i.e., those of the RBL and EBL signals and of any mixture of the two). The model also yields continuous predictions of how motor variability decreases as learning shifts from purely RBL signals (*β* = 0) towards EBL (higher *β*; Fig. 2c).

#### Target generalisation

Following Izawa and Shadmehr ^2^, we also measured generalisation by testing how much the policy was able to generalize the learned rotation perturbation (i.e., in the previous task) to novel target locations. The roation perturbation was learned for a target place at a 0-degree angle from the starting point and generalisation was tested for targets placed between −30 and +30 degress from the starting point, as in Izawa and Shadmehr ^2^ (see Methods for more details). Our model predicts that the dopamine-driven RBL condition yields poorer generalisation to novel targets than the cerebellar-driven EBL condition (Fig. 2e), while mixing the two signals produces intermediate performance. This same pattern is observed in humans across the three conditions ^2^. The poorer generalisation under RBL arises because, during training, RBL converges to a worse policy, in terms of training error, than EBL or the Mixed condition (Fig. 2g). Finally, to understand why humans rely less on cerebellar-driven EBL when sensory feedback is degraded ^2,30,36,37^, we measured the ℓ_2_-norm between the ground-truth gradient and the estimated EBL gradient across sensory-noise levels. The EBL gradient becomes increasingly inaccurate as sensory noise grows (Fig. 2l), impairing performance (Fig. 2k) e.g., ^30^. We further explore the link between sensory uncertainty and EBL, RBL combinations in the supplementary material.

#### Reward prediction errors drive motor variability changes

Finally, we examine another well-established effect in the motor neuroscience literature. During RBL, human motor variability increases following negative reward prediction errors and decreases following positive ones, efficiently regulating exploratory behaviour ^28,29^. Our action-gradient model reproduces this effect: fixing reward prediction errors to be consistently negative or positive and setting *β* = 0 (RBL only), the model predicts action variability to rise after negative errors and fall after positive ones (Fig. 2h). This follows directly from how the RBL action gradient modulates action variance in response to the sign of the reward prediction error.

### Action-gradient framework predict cerebello-basal ganglia interactions over learning

Here we show how our action gradient framework can explain well known cerebellar and basal ganglia motor learning impairments while offering novel predictions for future experimental validation. Following classical studies of cerebellar and basal ganglia motor learning, we test the framework on a set of motor learning tasks ^52,53,60–63^.

#### Visuomotor rotation task

We first evaluate the model on a classical cerebellar-dependent adaptation task in which subjects adjust to a sudden visuomotor rotation during endpoint arm-reaching movements (see Fig. S1) e.g., ^10,53,64,65^. We follow the experimental design of Tseng et al. ^52^, comparing healthy controls with cerebellar (CB) patients.

The model successfully reproduces the adaptation differences between control participants and CB patients (Fig. 3a), doing so solely through the distinct contributions of RBL and EBL action gradient estimates to learning. Healthy controls are best captured when learning is fully driven by the EBL action gradient—associated with cerebellar processing—resulting in lower residual errors (*β* = 1 in Eq. 2). In contrast, CB patients are best described when learning relies entirely on the RBL action gradient—associated with dopaminergic signaling (*β* = 0).

**Figure 3.**
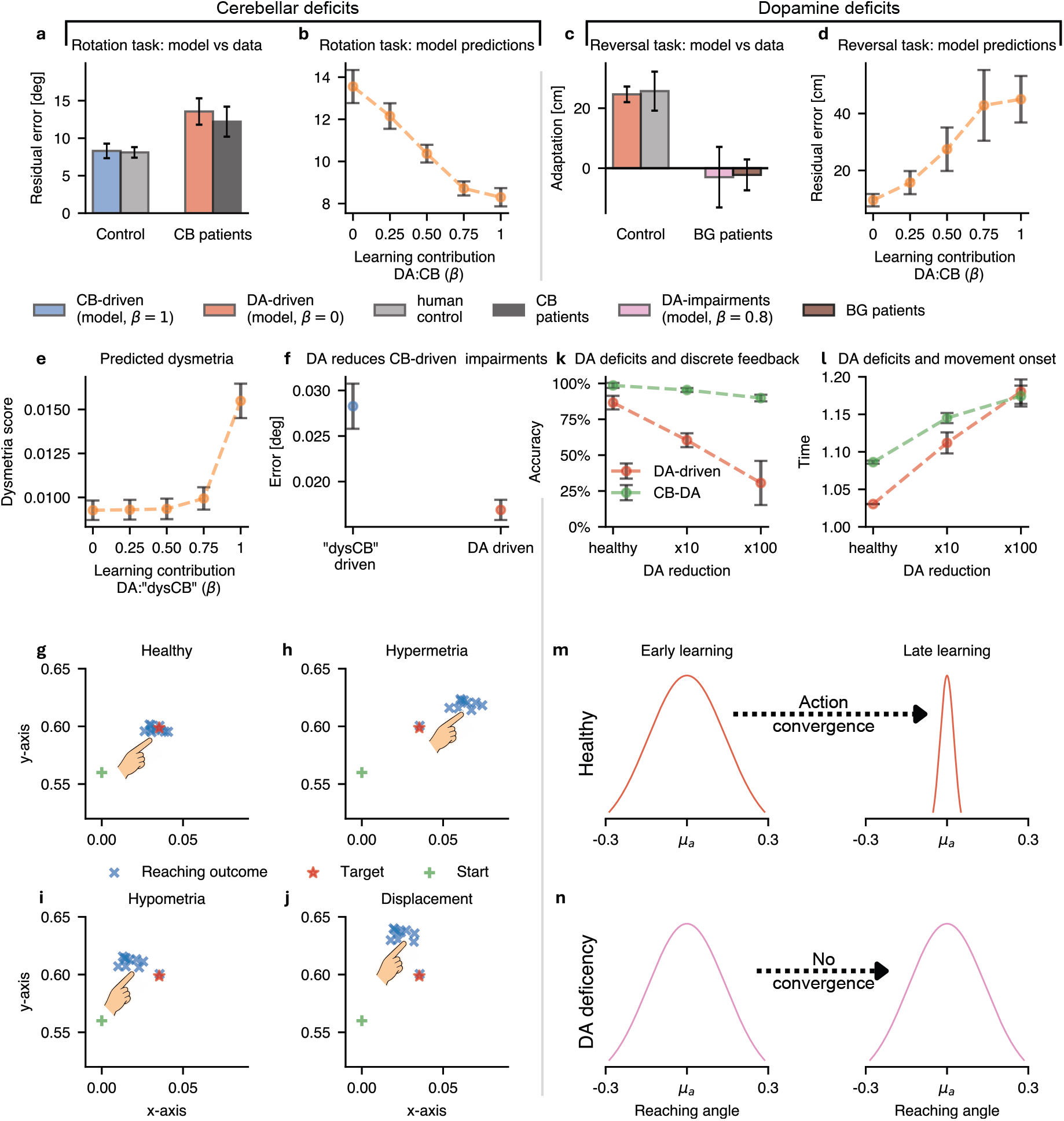
The action-gradient learning framework explains the impact of dopamine loss and cerebellar deficits on motor learning, while making predictions on their interactions. **a**, The framework replicates cerebellar patient deficits on a visuo-motor rotation arm-reaching task when adaptation receives no contribution from the cerebellar-driven EBL action gradient (i.e., consistent with cerebellar deficits). **b**, Adaptation to the visuomotor rotation task improves progressively as the contribution of the cerebellar-driven EBL action gradient increases relative to the dopamine-driven RBL action gradient. **c**, The framework replicates Basal Ganglia patient deficits on a visuomotor reversal arm-reaching task when adaptation receives a reduced contribution from the dopamine-driven RBL action gradient (i.e., *β* = 0.8), reflecting dopamine impairments. **d**, Adaptation to the visuomotor reversal task improves progressively as the contribution of the dopamine-driven RBL action gradient increases relative to the cerebellar-driven EBL action gradient. **e**, When the sing of the cerebellar-predicted sensitivity derivatives is systematically altered (i.e., dysCB), a more pronounced CB contribution to learning lead to increased Dysmetria symptoms during targeted arm reaching. **f**, Compensatory role of DA to erroneous cerebellar predictions (i.e., dysCB) driving lower reaching errors in a argeted arm reaching task. **g–j**, Distinct dysmetria symptoms emerge under the framework, including h) hypermetria, i) hypometria, and j) displacement, each corresponding to perturbing (i.e., flipping the sign of) a different component of the cerebellar predicted sensitivity derivatives. **k**, Dopamine deficiencies (10x or 100x reductions in dopamine levels) are associated with increased errors in a discrete feedback arm-reaching task (red), with performance partially recovering once sensory feedback is introduced, enabling a compensatory cerebellar contribution to learning (green, CB-DA). **l**, Movement onset time increases in proportion to the magnitude of dopamine deficiency (i.e., 10x vs. 100x), and the added cerebellar contribution fails to improve onset time or recover performance in this case. **m**, Healthy dopamine levels drive action convergence during learning, resulting in faster movement onset once learning is complete. **n**, Dopamine deficiencies prevent action convergence during learning within the framework. Error bars are computed based on performance across five random seeds. DA: dopamine, CB: cerebellum, dysCB: dysfunctional cerebellar predictions by randomly altering the sign of the cerbellar predicted sensitivity derivatives.

This dissociation suggests that, consistent with our theoretical framework, cerebellar impairment forces reliance on the suboptimal dopaminergic RBL gradient, leading to increased residual errors. Notably, in this task full visual feedback is available, which renders the RBL gradient less efficient than the EBL gradient. At the same time, the results indicate that CB patients retain a capacity for learning—albeit suboptimal—by leveraging the dopaminergic RBL pathway, in line with prior empirical findings e.g., ^38,66^.

The model further predicts how adaptation varies with the relative contribution of the cerebellar EBL and dopaminergic RBL gradients. Specifically, as the cerebellar contribution increases (i.e., as *β* increases), the residual error decreases monotonically, reaching its minimum when learning is driven entirely by the EBL gradient (Fig. 3b).

This prediction offers a potential explanation for the positive relationship between cerebellar symptom severity and adaptation errors reported by Tseng et al. ^52^. Patients with milder cerebellar impairments may retain partial access to the cerebellar EBL gradient, allowing them to outperform patients who must rely predominantly on the dopaminergic RBL gradient.

#### Visuomotor reversal task

Next, we show that the model reproduces adaptation in healthy controls and basal ganglia patients (Parkinson’s and Huntington’s disease) performing a visuomotor reversal task based on Gutierrez-Garralda et al. ^62^ (see Fig. S1). Unlike the rotation task, a reversal task inverts the visual feedback, making sensory error minimization counterproductive and causing error-based learning to degrade performance over training e.g., ^35,67,68^. Instead, successful adaptation is known to rely on reward-based learning (RBL) rather than error-based learning (EBL) e.g., ^27,62,63^. The reversal task therefore provides a natural complement to the rotation task, in which cerebellar EBL constitutes the optimal learning strategy.

The model reproduces the successful adaptation of healthy controls ^62^ when learning is driven entirely by the dopaminergic RBL action gradient (*β* = 0; Fig. 3c). In contrast, we found the (impaired) performance of basal ganglia patients were best reproduced by the model assuming a reduced contribution of the dopaminergic RBL action gradient to learning (i.e., *β* = 0.8 vs *β* = 0.0 in healthy) (pink). This finding is consistent with the well-established loss of dopaminergic signaling in Parkinson’s disease ^69–71^ and in later-stage Huntington’s disease ^72^, which in our model manifests as a diminished RBL action-gradient contribution to learning.

Finally, the model predicts that residual errors should increase as dopaminergic function declines, and therefore should correlate positively with symptom severity in reversal tasks (Fig. 3d). Although this prediction has not, to our knowledge, been tested directly, it is consistent with the broader observation that motor deficits worsen with disease progression in both Parkinson’s and Huntington’s disease ^73,74^.

#### Dysfunctional cerebellar predictions during reaching

In the visuomotor rotation task above, cerebellar impairment was modeled as a complete loss of the EBL action gradient. Nevertheless, the model also predicts the consequences of a different form of dysfunction, in which the EBL gradient remains available but is computed from erroneous cerebellar predictions—that is, from inaccurate sensitivity-derivative estimates 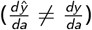. Such a scenario could arise when the cerebellum remains active but transmits incorrect predictions. We refer to this condition as “dysCB” (i.e., dysfunctional cerebellum) and investigate it in a standard reaching task, in which the model reaches to a target using a classical kinematic arm model under otherwise normal conditions ^75^.

Systematically perturbing the sign of the cerebellar-predicted sensitivity derivatives 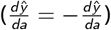 produced distinct and highly specific motor deficits (Fig. 3e–j). Specifically, the task required the cerebellum to predict four distinct sensitivity derivatives (i.e., the task had a 2D action space and a 2D output space, 2 *×* 2 = 4; see Fig. S2 in the Supplementary Material for a visual explanation). Strikingly, flipping the sign of each of these four cerebellar-predicted sensitivity derivatives individually reproduced a distinct, well-known cerebellar deficit: three of the four perturbations each mapped onto a classic clinical sign (h) hypermetria, (i) hypometria, and (j) displacement ^61,76,77^, indicating that individual components of the cerebellar sensitivity-derivative prediction correspond to specific, dissociable aspects of motor control. Interestingly, the remaining one sensitivity derivative perturbation had no effect on motor performance. When all four sensitivity derivatives were perturbed simultaneously, reaching accuracy deteriorated, producing a combined deficit that reflected a mixture of these symptoms (e.g., a blend of hypermetria, hypometria, and displacement) (Fig. 3f).

The model further predicts that this combination of symptoms should improve as learning shifts away from the dysfunctional cerebellar EBL gradient and toward the dopaminergic RBL gradient, resulting in lower dysmetria scores and reaching errors (Fig. 3e-f). This suggests that dysmetria arising from inaccurate cerebellar predictions may be mitigated by deliberately engaging reward-based learning. Although this prediction remains to be tested directly, it is consistent with emerging evidence that reinforcement-based training can improve motor performance in cerebellar patients, highlighting a potential avenue for rehabilitation based on reward feedback e.g., ^66^.

#### Dopamine deficiency, action uncertainty and slow movement onset

A cardinal feature of Parkinson’s disease is a generalized slowing of movement bradykinesia, ^78^, accompanied by prolonged decision-making times ^79^. Both are thought to arise, at least in part, from the widespread dopaminergic (DA) deficits associated with the disease ^69–71^. Here we ask whether our action-gradient framework can provide a mechanistic account of this relationship. Specifically, we investigate how reductions in dopamine alter the uncertainty of the learned action distribution, which in turn is predicted to determine movement onset time higher action uncertainty delays movement initiation, ^80–82^.

To relate action uncertainty to movement onset, we couple our framework to a standard single-boundary drift-diffusion model ^83^, in which decision time depends on uncertainty over the target action (see Methods). We test this in a classical reward-based endpoint-reaching task in which learning is driven exclusively by binary success/failure reward feedback, with no sensory feedback available ^39^(as in Fig. 2a). This paradigm aims to isolate the contribution of the dopamine-dependent RBL action gradient, because the cerebellar EBL action gradient cannot be computed in the absence of sensory feedback e.g., ^2,19,84^. We compare three conditions: a healthy baseline with normal dopamine levels, and two dopamine-deficient conditions in which reward prediction errors (δ_*t*_) are uniformly scaled down by factors of 10 and 100, respectively (i.e., mimicking a reduction dopamine release in repose to a reward prediction error).

The model predicts that movement onset time increases monotonically with dopamine deficiency (Fig. 3l). This is because under healthy dopamine levels, learning substantially reduces uncertainty over the target action, enabling rapid movement initiation (Fig. 3l,m). This follows directly from the RBL action gradient update, which reduces the uncertainty of the policy as learning progresses towards the optimal policy. However, as dopamine levels decline, action uncertainty fails to decrease, maintaining high action uncertainty through learning, which then drives delayed movement onset (Fig. 3n). These results suggest a potential computational link between dopamine deficiency, elevated action uncertainty, and bradykinesia ^85^. Interestingly, Fig. 3l shows that introducing visual feedback does not allow the cerebellum to compensate for the increased movement onset time caused by reduced DA availability (CBDA, green): onset time remains elevated even with cerebellar contribution. Furthermore, we find that DA deficiency degrades not only movement onset time but also reaching accuracy (Fig. 3k, red). In this case, however, allowing the cerebellum to contribute to learning by introducing sensory feedback (i.e., via the EBL action gradient) leads to partial compensation in motor accuracy (Fig. 3k, green). This finding offers a mechanistic explanation for the well-established observation that Parkinson’s patients benefit disproportionately from explicit sensory cues during movement ^86,87^: sensory feedback enables cerebellar error-based learning to compensate, at least partially, for impaired dopaminergic reinforcement learning.

### Cerebellar-driven network accounts for interference and savings

Savings and interference are two canonical phenomena of motor learning that have been extensively documented in visuomotor adaptation e.g., ^88–91^. Savings refers to the accelerated relearning of a previously experienced perturbation, indicating that the nervous system retains a memory of the earlier adaptation. Interference refers to the loss of this benefit when an opposing perturbation is learned between two exposures to the original perturbation, suggesting that the intervening adaptation disrupts retrieval of the initial motor memory.

Here, we demonstrate that our action-gradient framework provides an account of both phenomena through the cerebellar EBL pathway. In the framework, the cerebellum learns the sensitivity derivatives (the action-to-outcome mapping, 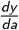 required to compute the EBL action gradient. Once these derivatives have been learned for a given perturbation (e.g., a clockwise 30° visuomotor rotation), they can be retrieved when the same perturbation reappears, allowing the model to adapt more rapidly (savings). Conversely, if an opposing perturbation (e.g., a counter-clockwise 30° rotation) is experienced before re-exposure, learning the new sensitivity derivatives may overwrite the previously learned ones. As a result, the original derivatives are no longer available when the first perturbation returns, preventing accelerated relearning (interference).

This account relies on one key assumption: the cerebellum must distinguish the perturbed context from the unperturbed baseline in order to retrieve the appropriate sensitivity derivatives for each. Such context-dependent switching of cerebellar representations has recently gained both theoretical and experimental support ^50,92–96^.

To model this contextual switching, we augment the cerebellar network with context-specific inputs, following the approach of Pemberton et al. ^50^. Separate cerebellar input pathways encode different task contexts, enabling the network to learn and retrieve context-dependent sensitivity derivatives (see illustration in Fig. S2 and details in Methods). We evaluate the resulting model on two classical visuomotor adaptation paradigms ^88,89^, which probe savings and interference through repeated visuomotor rotations during endpoint reaching.

In the savings paradigm, subjects first adapt to a 30° visuomotor rotation and, following a washout period with veridical feedback, are re-exposed to the same rotation (Fig. 4a). The model closely reproduces the experimental findings ^88^, exhibiting faster adaptation during the second exposure than during the first (Fig. 4d). Examining the cerebellar sensitivity-derivative predictions throughout the task (Fig. 4b,e,h,k) reveals a large update when the perturbation is first introduced (first yellow region), but only a minor adjustment when it reappears (second yellow region). This pattern indicates that the cerebellum retrieves the previously learned sensitivity derivatives, enabling faster relearning.

**Figure 4.**
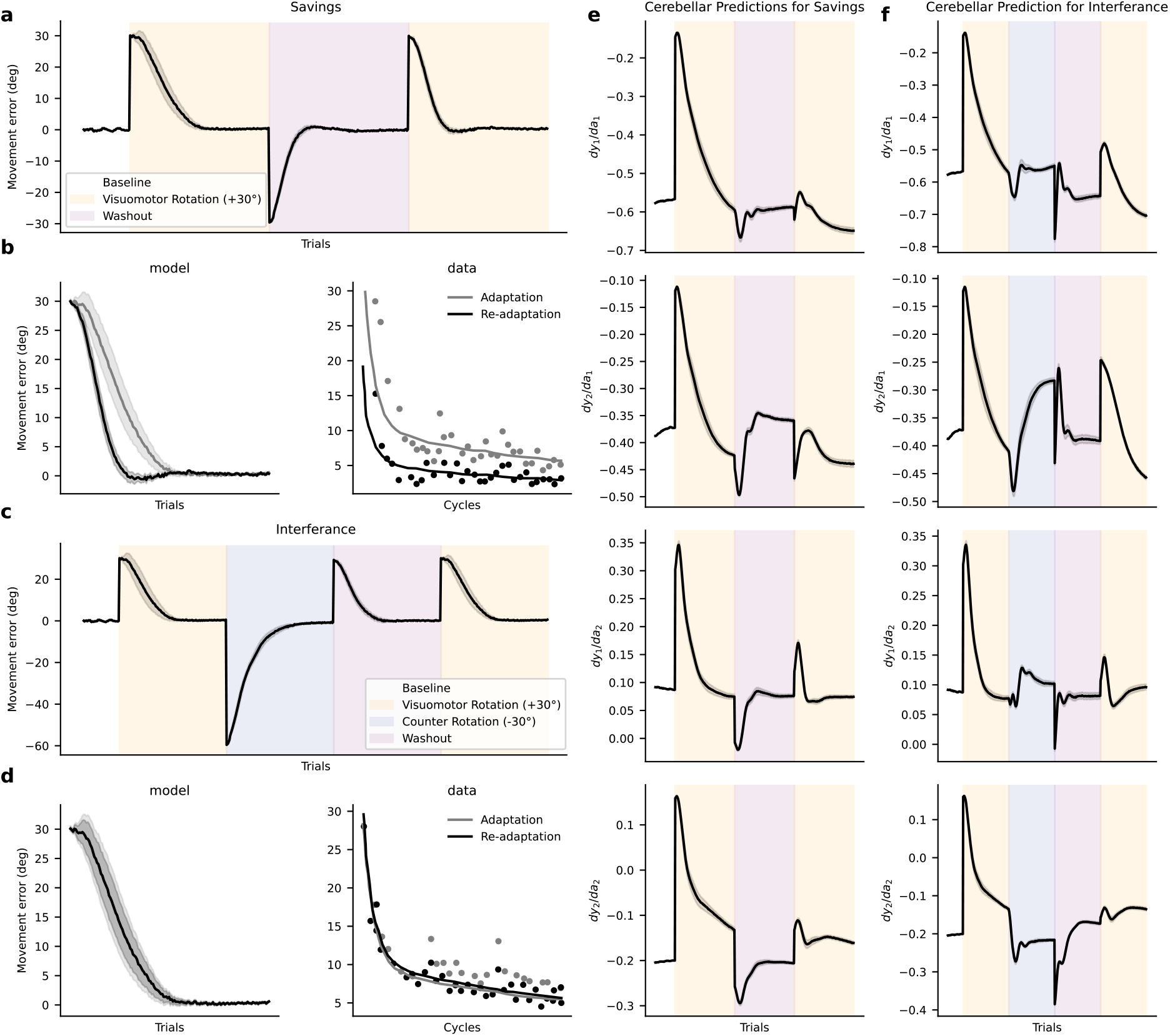
Cerebellar-driven network accounts for interference and savings. **a**, Model-predicted adaptation in the *savings* paradigm. An initial +30° visuomotor rotation (yellow) is introduced, followed by a washout period with veridical feedback (pink), after which the same +30° rotation is reintroduced (yellow). **b,e,h,k**, Cerebellar sensitivity-derivative predictions 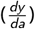 during *savings* (see Fig. S2 for the model schematic and the four sensitivity derivatives used in the reaching task). **d**, Comparison of adaptation in the action-gradient learning framework (left) and the experimental data of Krakauer et al. ^88^ (right), demonstrating *savings*. **g**, Model-predicted adaptation in the *interference* paradigm. An initial +30° visuomotor rotation (yellow) is followed by an opposing −30° rotation (purple), a washout period with veridical feedback (pink), and finally re-exposure to the original +30° rotation (yellow). **c,f,i,l**, Cerebellar sensitivity-derivative predictions 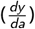 during *interference* (cf. Fig. S2). **j**, Comparison of adaptation in the action-gradient learning framework (left) and the experimental data of Krakauer et al. ^88^ (right), demonstrating *interference*.

In the interference paradigm, an equal-magnitude but opposite rotation (−30°) is inserted between the first and second +30° rotations (Fig. 4g). Again, the model reproduces the experimental findings ^88^: adaptation to the second +30° rotation is no longer faster than adaptation to the first (Fig. 4j). In contrast to the savings paradigm, the cerebellar sensitivity-derivative predictions (Fig. 4c,f,i,l) undergo substantial updating when the second +30° rotation is encountered (second yellow region), indicating that the previously learned derivatives were washout. Within our framework, this occurs because learning the intervening −30° rotation overwrites the sensitivity derivatives of the initial rotation, thereby abolishing the savings effect and producing interference. In the supplementary material, we further show that savings arise from the cerebellum cashing the sensitivity derivative in a context-dependent manner e.g., ^50,92–96^, since this phenomenon does not occur when learning is driven solely by the dopaminergic RBL action gradient (see Fig. S7, i.e., *β* = 0 in our model).

### Cerebellum controls the sign of dorsal striatal plasticity

Thus far, we have evaluated our unified RBL–EBL framework against classic motor learning paradigms. Here, we leverage the model to derive novel, testable predictions that can guide future experimental work and provide targets for model validation. In our system-level implementation, the cerebellum predicts the sensitivity derivative component of the EBL action gradient (i.e., 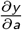 in Eq. 2). This component instructs how dorsal striatal synapses encoding actions must adapt to reduce sensory errors *e*, thereby driving striatal EBL (Fig. S2). Crucially, theoretical work demonstrates that successful EBL requires only the correct sign of 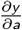, whereas its magnitude merely scales the rate of learning e.g., ^27,46^. A central prediction of our framework is therefore that cerebello-dorsal-striatal projections ^21,25^ should primarily dictate the direction of dorsal striatal plasticity (i.e., long-term potentiation vs. depression) in response to cortical sensory-error signals.

Fig. 5 illustrates this mechanism for both a fixed positive (Fig. 5a) and a fixed negative (Fig. 5b) cortical sensory error. In each case, the cortical sensory error is held constant while we artificially and periodically flip the sign of the cerebellar-predicted sensitivity derivative every few trials. This periodic sign-flip triggers an immediate, corresponding switch between LTP and LTD (or vice-versa) at dorsal striatal synapses. This illustrates across both error conditions, the sign of the cerebellar-predicted sensitivity derivative control the direction of dorsal striatal plasticity, in relation to the (cortical) sensory error.

**Figure 5.**
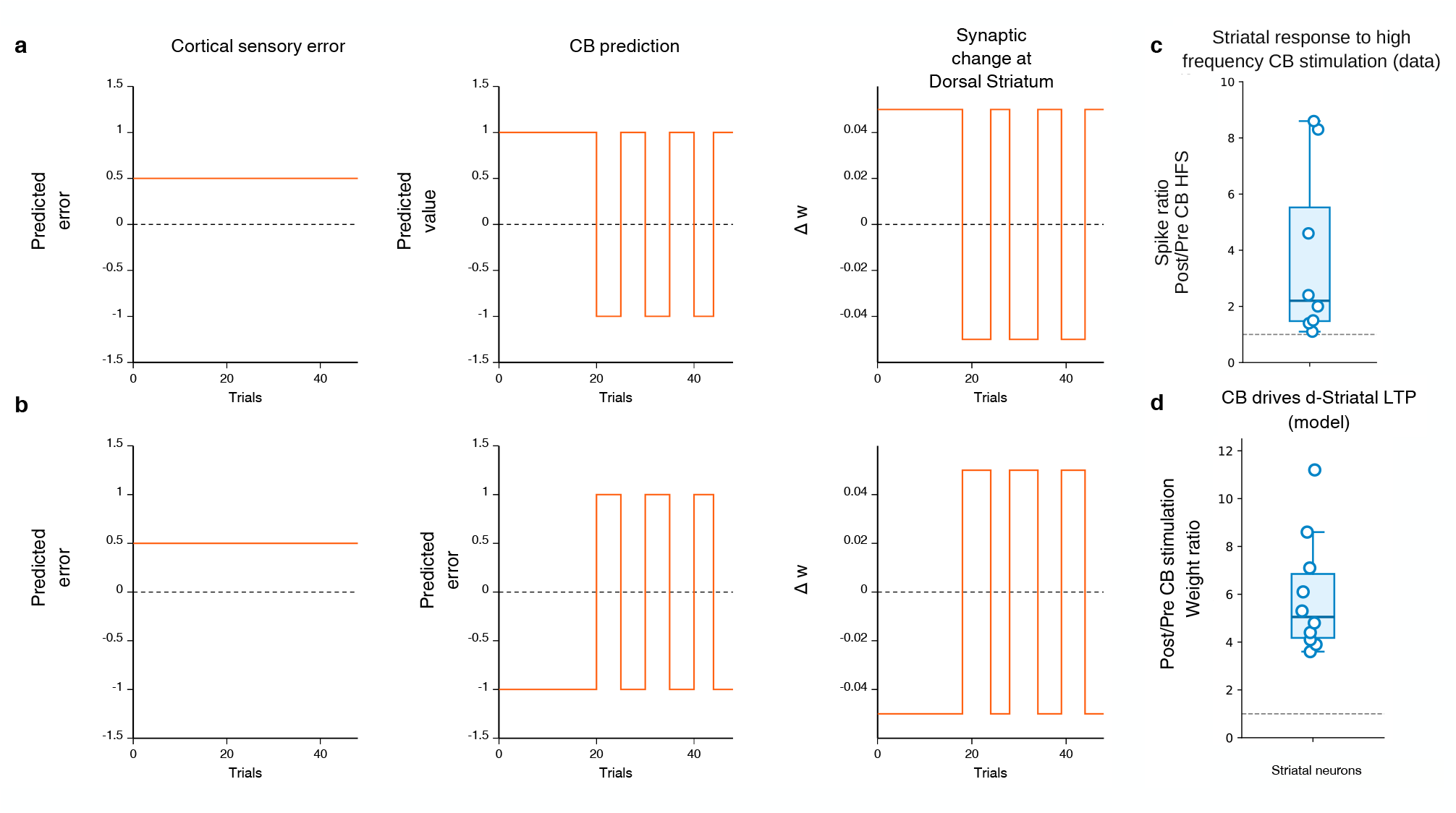
Cerebellar predictions controls the sign of dorsal striatal plasticity. **a,b**, Policy (e.g., dorsal striatum) weight changes (Δ*w*) under fixed positive 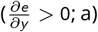 or negative 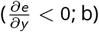 (cortical) sensory errors across trials. Artificially and periodically flipping the sign of the cerebellar-predicted sensitivity derivative 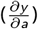 triggers an immediate, corresponding switch between LTP and LTD (or vice-versa) at dorsal striatal synapses. **c**, Experimental demonstration of dorsal striatal LTP induced by high-frequency stimulation (HFS) of the cerebello-thalamo-striatal pathway (adapted from Chen et al. ^24^). **d**, Simulated LTP across 10 randomly initialized policy weights when cerebellar and sensory inputs relay positive sensitivity derivatives 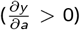 and positive sensory errors 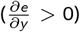, to the dorsal striatum (policy) respectively

While this prediction warrants direct empirical validation, it is strikingly consistent with work by Chen et al. ^24^, who demonstrated that high-frequency cerebellar stimulation in mice converts dorsal striatal plasticity from LTD to LTP exclusively in striatal neurons recipient of cerebellar input. Crucially, this plasticity switch occurs at short latency—a feature essential for trial-to-trial motor adaptation. Within our framework, such rapid subcortical convergence of cerebellar sensitivity derivatives and cortical sensory errors provides the exact signal required to compute and apply the EBL action gradient alongside dopaminergic reward signals.

To test the plausibility of this mapping, we confirmed that our model qualitatively reproduces the physiological findings of Chen et al. ^24^, showing robust LTP at policy-encoding synapses under joint cerebellar and cortical activation (Fig. 5c,d). Across 10 random weight initializations, setting the cerebellar and cortical inputs to encode positive sensitivity derivatives 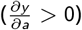 and positive sensory errors (*e >* 0), respectively, drove consistent synaptic potentiation. This mapping assumes that the high-frequency stimulation used by Chen et al. ^24^ translates to a positive sensitivity derivative in our model, as our framework does not explicitly model spiking rates or stimulation protocols. Interestingly, a comparable subcortical convergence of cerebellar error cues and basal ganglia reward signals has been proposed to account for song learning in songbirds ^26^.

### Dopamine-driven RBL drives memory consolidation

The reinforcement learning literature demonstrates that the RBL weight update (Eq. 1) extends to off-policy settings ^97^, where updates are computed on past experiences replayed from a memory buffer. A prediction of our action gradient framework is that the brain would use similar dopamine-driven action-gradients during offline memory consolidation. We demonstrate a similar outcome by showing that motor performance acquired during a reward-based task can be consolidated *post-learning* by applying the dopamine-driven RBL update during replay of stored experiences.

To test this principle, we use the endpoint reaching task described above, in which a policy learns actions to maximize a binary reward (Fig. 2a). To model post-learning retention, we introduce a standard synaptic weight decay in the policy network (see Methods) and probe performance at regular intervals over an extended retention period, aligning with classic reward-based motor learning paradigms ^39^. Our framework predicts a substantial boost in long-term retention when dopamine-driven offline replay occurs during this period (Fig. 6a,c, solid vs. light red line). We next examine how dopamine availability during retention modulates this consolidation gain. Following the classic reward-prediction error hypothesis ^4,42^, we model dopamine deficiency by scaling down the reward prediction error δ that drives the offline RBL update by one or two orders of magnitude (as in the previous task). Under RPE/dopamine depletion, the framework predicts a steep decline in retention due to impaired offline RBL consolidation (Fig. 6d). This computational mechanism provides a potential explanation for why Parkinson’s disease patients often exhibit impaired retention of motor skills ^98–100^: while neural replay of past experiences may remain intact, insufficient dopaminergic signaling degrades the synaptic updates required for offline consolidation. Finally, we explore how our action gradient framework extends reward-driven consolidation to error-based learning (EBL). The framework posits that reward-driven RBL and cerebellar-driven EBL signals converge within the same policy-encoding neurons (e.g., in the dorsal striatum) rather than driving independent policies. A key consequence of this shared architecture is that whenever dopamine-driven RBL acts offline to consolidate motor memories, it inherently consolidates motor functions originally acquired via cerebellar EBL. Indeed, after training the task using EBL, we demonstrate that memory consolidation can occur solely through reward-driven RBL updates operating on replayed experiences (Fig. 6b,c, blue). This prediction provides a potential explanation for why combining reward feedback with sensory error signals enhances long-term motor retention ^18,31–34^. It is important to stress that even during CB-driven learning, reward signals may still be available (e.g., subjective evaluations of the utility or success of sensory outcomes), which can then be leveraged for offline memory consolidation.

**Figure 6.**
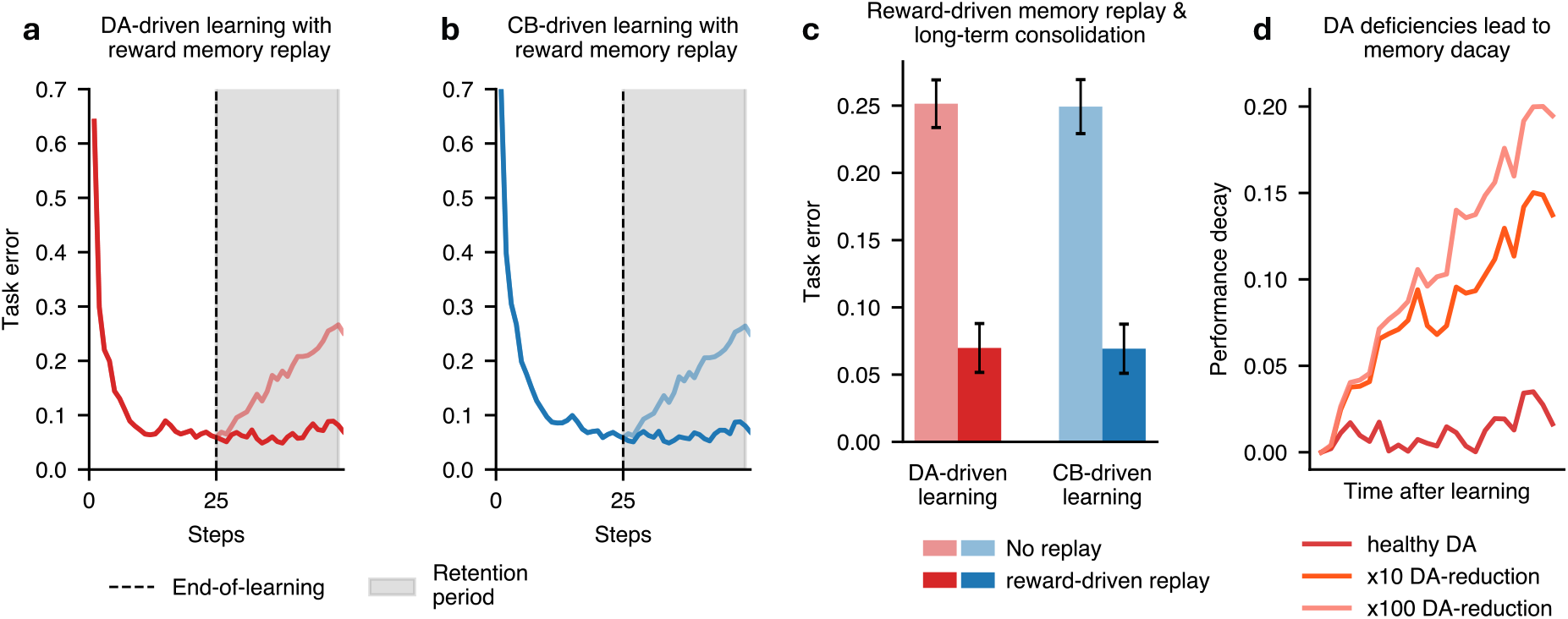
Replay of rewarding events can support consolidation in both error-based and reward-based learning settings. **a-b**, Post-learning reward-driven memory replay (gray area) prevents performance decays of both DA-(red) and cerebellar- (blue) driven learned motor functions. **c**, Post-learning reward-driven (DA-driven) memory replay drives task consolidation of both (*β* = 0) DA- (red) and (*β* = 1) cerebellar- (blue) driven learned motor functions. **d**, DA deficiencies ( x10 or x100 reductions in DA levels) are predicted to impair memory consolidation resulting in poorer motor performances as more time elapses from the end of the task. DA: dopamine, CB: cerebellum, rwd: reward.

### Optimal combination of CB and DA-dependent learning depends on task expertise

A fundamental prediction of our action gradient learning framework is that the brain relies on a (*β*-weighted) combination of the cerebellar-driven EBL and dopamine-driven RBL teaching signals (the action gradients). The optimal weighting of the two should be influenced by the accuracy of the cerebellar sensitivity-derivative estimate, which represents a key component for accurate EBL (Fig. 7a). Crucially, these sensitivity derivatives must themselves be learned ^27,46^. A novice tennis player, for instance, does not yet know how small changes in racket force or angle affect the ball’s trajectory; this sensitivity improves only with expertise. A novice’s cerebellar estimates of the sensitivity derivative are therefore unreliable, yielding inaccurate EBL action gradients. In this case, we expect optimal learning should be driven by the dopamine-driven RBL signal, which needs no prior knowledge of the action-to-outcome mapping. With practice, the cerebellum learns to estimate the sensitivity derivatives more accurately, enabling accurate EBL signals and promoting faster and more precise learning. Therefore, our action gradient learning framework predicts that optimal learning should involve a shift in the relative contributions of RBL and EBL across learning stages: novices should rely more heavily on RBL, whereas with practice they should gradually transition toward greater reliance on EBL.

**Figure 7.**
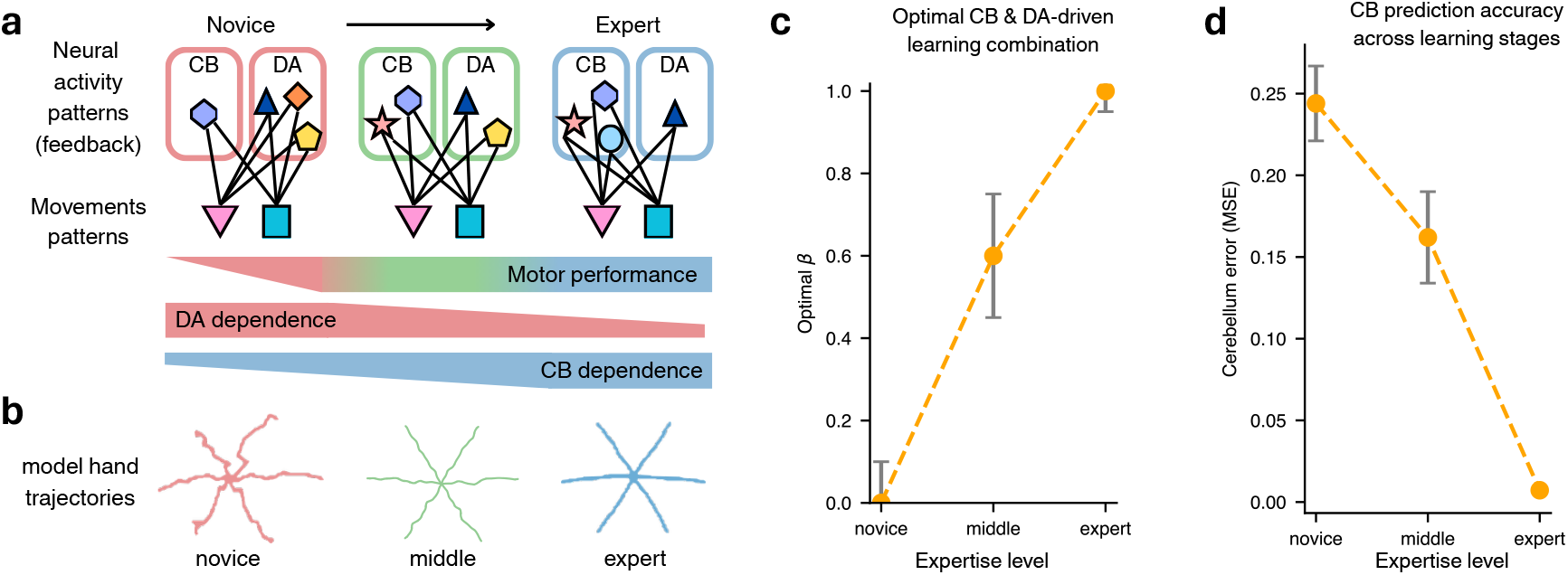
Optimal combination of CB and DA-dependent learning depends on task expertise. **a**, Schematic illustrating long-timescale shifts in CB and DA engagement during motor learning, highlighting their necessity and contributions to movement expression across stages. **b**, Comparison of the model-generated hand drawing trajectories for novice, intermediate, and expert task levels. **c**, Our framework predicts the optimal combination of cerebellar-driven EBL and DA-driven RBL learning signal (as determined by *β* in Eq. 2) depends on the level of task expertise. **d**, The resulting cerebellar network error as a function of task expertise.

Testing this prediction requires studying learning across all of its stages—from complete novice to expert—which calls for a de novo learning paradigm (i.e., learning a new task from scratch) rather than the adaptation paradigms used above, where the model already begins with a well-learned baseline policy and only needs to adjust it. To this end, and inspired by classical motor learning studies, we designed a task where the policy must learn to hand draw one of six straight-line trajectories entirely from scratch (Fig. 7b). Each line is drawn over multiple timesteps based on a cue presented only at the first step. To generate the correct trajectory, the policy—a recurrent neural network (RNN)—must maintain this cue information across all timesteps (see Methods), closely following previous work ^48–50^.

Fig. 7c,d show that the optimal balance between the EBL and RBL action gradients is driven by the accuracy of the cerebellar prediction across learning stages. Early in learning (novice phase), the cerebellar estimate of the sensitivity derivative carries a large error (Fig. 7d), making the optimal strategy to rely entirely on the dopamine-driven RBL action gradient (*β* = 0; Fig. 7b). As the cerebellar error decreases (intermediate phase), optimal performance shifts to a balanced mixture of both signals (*β* ≈ 0.6; Fig. 7c,d). Finally, once the cerebellar error reaches zero (expert phase), the optimal strategy relies exclusively on the EBL signal (*β* = 1; Fig. 7c,d; cf. Fig. 7a).

We speculate that in real-world motor control, cerebellar error rarely reaches zero because environmental and bodily dynamics change continuously (e.g., wind, muscle fatigue/variability). As a result, the RBL action gradient likely maintains a persistent contribution even in highly skilled individuals. Beyond expertise, other factors also constrain the accuracy of cerebellar sensitivity derivatives and, by extension, the optimal EBL–RBL balance. Consistent with this, we demonstrate in the Supplementary Material (Fig. S4) that sensory noise similarly shapes this trade-off: higher noise levels degrade the accuracy of the EBL gradient, shifting the optimal strategy toward greater RBL reliance. Together, these results indicate that adaptive behavior in natural environments demands dynamic integration of RBL and EBL signals—governed by *β* as predicted by our action gradient framework.

## Discussion

Here, we introduced a novel computational framework demonstrating how the brain integrates sensory error-based learning (EBL) and reward-based learning (RBL) by converting both teaching signals into a shared action-gradient representation that drives a common downstream policy.

Mapping this framework to a systems-level circuit provides a mechanistic foundation for the growing evidence of functional interactions between the cerebellum and basal ganglia ^15,21–24,26^. The model accounts for classical findings across both domains ^52,61,62,77,88^ while generating distinct, testable predictions about their joint contributions to motor adaptation.

Specifically, the model predicts that the policy-encoding dorsal striatum learns not only from dopaminergic reward prediction errors ^5,6,8^, but also from cerebellar feedback conveying EBL action gradients. Consistent with this fast, trial-by-trial mechanism, Chen et al. ^24^ demonstrated that cerebellar stimulation rapidly modulates dorsal striatal plasticity. Aligning with these findings, our model reproduces the specific synaptic LTP patterns reported by Chen et al. ^24^ during paired cerebellar and cortical stimulation. Furthermore, perturbing cerebellar predictions within the model yields classic dysmetria (hypermetria and hypometria) ^61,76,77^, while simulating dopamine loss reproduces characteristic basal ganglia deficits such as bradykinesia and elevated motor variability ^78,79^. Together, this framework offers a unified computational account of motor control and pathology across these two traditionally distinct systems.

Our central contribution is the proposal that cerebellar- and dopamine-driven teaching signals share a common action-gradient representation, allowing them to directly govern synaptic plasticity within a single downstream policy. This departs fundamentally from previous models of cerebellar–basal ganglia interactions, which typically assume that error-based (EBL) and reward-based learning (RBL) update separate, distinct policies—such as one in the cerebellum and another in the basal ganglia ^2,8,16,17^. This distinction is crucial: dual-policy accounts lack a clear mechanism for coordinating reward and error signals toward a unified behavioral goal. By expressing both teaching signals as action gradients, our framework provides a shared computational substrate that seamlessly integrates learning updates within a single policy (in the Supplementary Material, we detail why coordinating multiple independently updated policies poses a fundamental computational challenge).

Furthermore, integrating cerebellar EBL and dopamine-driven RBL within a shared action-gradient space provides two major computational advantages. First, it allows the relative contribution of each learning signal to adapt dynamically across skill acquisition. During *de novo* learning, cerebellar estimates of sensitivity derivatives are initially imprecise, making dopamine-driven RBL the primary, most reliable driver of updates. As task dynamics are learned, cerebellar predictions improve, and EBL action gradients become increasingly informative. At expert performance, highly accurate cerebellar predictions render EBL the dominant and most efficient mechanism for fine-tuning policy updates.

This dynamic shift reconciles the long-standing distinction between *de novo* motor learning – which exhibits signature features of reinforcement learning policy gradient methods ^101^ – and sensorimotor adaptation, which is dominated by EBL ^10,35^. Because standard adaptation paradigms perturb well-learned behaviors (such as reaching) where the cerebellum already possesses accurate forward models, EBL naturally predominates – not because the two processes rely on distinct systems, but because cerebellar action gradients are already highly reliable.

Second, a shared action-gradient space provides a straightforward mechanism for memory consolidation. Dopamine-driven RBL can drive offline consolidation via experience replay. By directing both cerebellar and dopaminergic signals to the same action-encoding synapses, our framework allows dopamine-dependent replay to consolidate motor memories regardless of whether they were initially acquired through sensory errors or rewards. Conversely, dual-policy architectures require separate, synchronized consolidation mechanisms to keep memories aligned across systems—a computationally demanding requirement that restricts knowledge transfer between learning regimes. This single-policy consolidation mechanism offers a computational rationale for why combining reward with sensory feedback significantly enhances long-term motor memory retention ^18,31–34^.

Our framework faces three principal limitations. First, it is formulated at Marr’s algorithmic level ^102^. While it maps abstract variables—like action gradients onto systems-level circuits, it does not specify the precise spiking, synaptic, and local circuit mechanisms that compute them. Second, the balance between EBL and RBL is governed by a static weighting parameter (*β*), leaving open the question of the brain dynamically adapt this arbitration across contexts.

Third, our EBL implementation focuses on task errors without modeling sensory prediction errors ^90,103^ or cognitive strategy re-aiming, both of which modulate motor adaptation.

Crucially, while we focus on cerebellar–basal ganglia interactions, this action-gradient framework is not restricted to the dorsal striatum. It applies broadly to any circuit integrating dopaminergic, cerebellar, and sensory inputs to encode actions. Downstream policies could thus be implemented in distinct neural substrates depending on the domain—such as motor cortex for complex motor behavior ^104–107^ and low-dimensional embeddings ^108^ or prefrontal cortex for cognitive control ^109^ – with cerebellar and dopaminergic signals providing the requisite EBL and RBL action gradients.

Consistent with this view, both cerebellar and dopaminergic projections modulate synaptic plasticity across multiple cortical areas. In primary motor cortex, dopaminergic signaling ^110–112^ and cerebellar outputs ^113–115^ both regulate synaptic plasticity. Similarly, converging evidence links dopaminergic ^116–118^ and cerebellar pathways ^119–121^ to prefrontal plasticity. Testing our framework’s predictions in these cortical circuits is an important venue for future research.

More broadly, understanding how the brain coordinates learning across interacting systems remains a fundamental challenge, as existing accounts remain largely fragmented. Our framework demonstrates that action gradients can serve as a common computational substrate, integrating error- and reward-based learning to update a shared policy. This framework bridges classical theories of motor adaptation based on directed policy learning ^35^ with reinforcement learning policy gradients ^122^, while accounting for decades of experimental findings across the cerebellar and basal ganglia literature. Rather than acting as competing systems, EBL and RBL serve as complementary teaching signals unified by a common action-gradient representation – providing a foundation for future experimental and theoretical work on multi-system learning.

## Acknowledgements

We would like to thank the Neural & Machine Learning group, John Krakauer, Juan Gallego and Manuel San Silvestre for useful feedback. MG was funded by the Wellcome Trust Transition Fellowship (S122871-115), CF was funded by a “La Caixa” Doctoral INPhINIT Retaining Fellowship (CCA 0404020203) and RPC by an EPSRC New Investigator Award (EP/X029336/1) and an ERC Research Starting Grant (via ERC-UKRI Frontier Guarantee EP/Y027841/1).

## Author contributions

The author contributions are as follows:

Conceptualisation: MG and RPC. Methodology: MG, CF, LA and RPC. Simulations: MG and CF. Investigation: MG, CF and RPC. Visualisation: MG, CF and RPC. Supervision: RPC. Writing: MG, CF, and RPC.

## Methods

### Reward and error based learning

Below we provide an overview of the two forms of learning that we consider here, which are the building blocks for our action-gradient model.

#### Reward-based Learning (RBL)

RBL is thought to drive synaptic plasticity of downstream motor areas (i.e., encoding the policy *π*_*ϕ*_) according to the following synaptic weight update i.e., the stochastic policy gradient updates in actor-critic algorithms, ^4–7^,

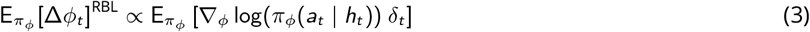

where Δ*ϕ*_*t*_ denotes the change in synaptic weight, while δ_*t*_ = *r*_*t*_ − *v*_*θ*_(*s*_*t*_) represents the reward prediction error following action *a*_*t*_ for some observed reward signal *r*_*t*_. Finally, *v*_*θ*_(*s*_*t*_) denotes the expected reward given some sensory cue/state, *s*_*t*_. This synaptic weight update assumes that reward prediction errors provide the learning signal to update the synaptic weight of the policy *ϕ*, increasing the probability of actions that resulted in positive reward prediction errors, while decreasing the probability of actions that resulted in negative reward prediction errors (i.e., moving towards highly rewarding actions and away from low-rewarding actions). Importantly, Garibbo et al. ^27^ show how we can derive the RBL action gradient in Eq. (1) from the standard RBL synaptic update above, assuming a Gaussian policy. Here, we report the derivation for the mean parameter *µ*_*ϕ*_ of the Gaussian policy (a similar derivation can be obtained for the standard deviation parameter *σ*_*ϕ*_),

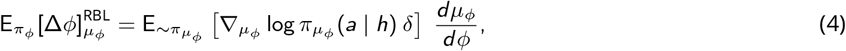

where we simply used the chain rule to separate the parameter updating term 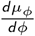 from the mean parameter *µ*_*ϕ*_ gradient (we dropped the subscript *t* for ease of reading). Next, we substitute the Gaussian policy in the expression,

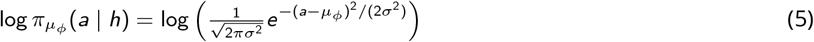

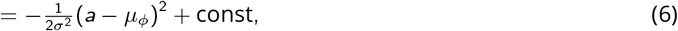

where the constant is independent of *µ*_*ϕ*_. Differentiating by *µ*_*ϕ*_,

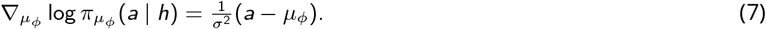

And substituting this into Eq. (4), we get,

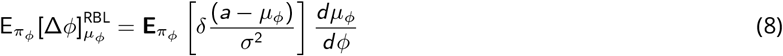

which is exactly the RBL synaptic weight update in Eq. (1).

In the brain, the RBL synaptic update has been related to the function of the Basal Ganglia. The policy, *π*_*ϕ*_, is associated with the dorsal striatum, which appears key for action selection ^5,123^. Conversely, the ventral striatum is thought to play the role of a critic, *v*_*θ*_, given its prominent role in value estimation (i.e., providing expected rewards) ^4,124^. Finally, striatal dopamine projections are thought to convey reward prediction errors, δ_*t*_, driving learning in both downstream dorsal (policy) and ventral (critic) striatum ^5^. This proposal is in line with the evidence of extensive dopaminergic projections to both striatal areas as well as striatal dopamine activity mimicking reward prediction errors (i.e., increasing in response to unexpected rewards and suppressing in face of unexpected lack of rewards) ^42,124,125^.

#### Error-based learning (EBL)

EBL is thought to drive synaptic plasticity of downstream motor areas (i.e., encoding the policy) according to the following synaptic weight update ^27,35,46,126^:

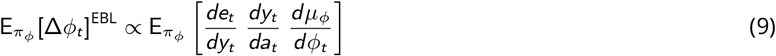

where Δ*ϕ*_*t*_ again denotes the change to the synaptic weights at trial *t*. The term *y*_*t*_ represents the sensory outcome in response to action *a*_*t*_, while *e*_*t*_ represents a (sensory) error signal following outcome *y*_*t*_. In motor learning paradigms, this error signal is typically computed as the squared distance between the final, *y*_*t*_, and the desired, *y*^∗^, (sensory) outcomes at any given trial *t* (i.e., *e*_*t*_ = (*y* (*a*_*t*_) − *y*^∗^)^2^) (e.g., the distance from the goal state after taking action *a*_*t*_). In Eq. (9), the operator 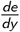 can be thought of as the ‘directed’ sensory error, encoding the direction to change the sensory outcome *y* to reduce the error *e* (e.g., signaling to throw more to the right to hit a target in a dart game). The operator 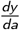 provides an action-to-outcome ‘interface’, enabling the mapping of (directed) sensory errors, 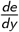, to motor changes (e.g., instructing how to change the action to throw more to the right). This quantity is commonly referred to as the sensitivity derivative ^46^, a term we adopt throughout. The sensitivity derivative is required because actions, *a*, and sensory outcomes, *y*, lie in different spaces (i.e., motor and sensory, respectively), necessitating an explicit mapping between them. Finally, 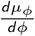 determines how to change the synaptic weights to change the (mean) action in the desired direction. In the brain, EBL has broadly been associated with the cerebellum (CB) ^9–11^.

### Policy representations

As mentioned in the Results, the actions (motor commands) are assumed to follow a Gaussian policy (reported here for the multidimensional case for completeness),

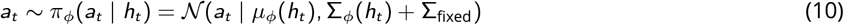

where *t* denotes the current trial, and the mean, *µ*_*ϕ*_, and (diagonal) covariance/variance, Σ_*ϕ*_, are parametrized by synaptic weights, *ϕ* (e.g., the synaptic weights of the area encoding the policy). We also assume a certain degree of motor variability to be fixed and irreducible (Σ_fixed_), since some degree of human motor variability is always present ^28^. These (policy) parameters, *ϕ*, are learned based on the action-gradient framework introduced in the “Action-gradient framework for reward- and error-based learning” section in the main text.

In visuomotor rotation, reversal, and discrete reward-based tasks, the policy parameterization consists of a linear model taking the desired target as input (*s* = *y*^∗^) and outputting the policy mean, *µ*_*ϕ*_, and variance, 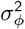 (i.e., for 1-dimensional actions):

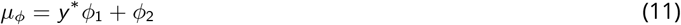

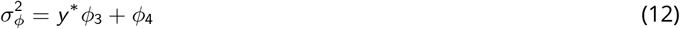

Note that in the experiments presented in the Supplementary Material with separate policies for EBL and RBL, we use two separate (linear) policies, where the RBL policy is exclusively updated based on RBL, while the EBL policy is exclusively updated based on EBL e.g., ^2,5,27,35^. Note also that this (linear) policy setup is standard practice in computational models of motor adaptation, where the action, *a*, typically represents the arm reaching angle e.g., ^2,27,35^.

In all the motor learning tasks involving the (non-linear) two-joint arm model, the (Gaussian) policy is parametrized by a 1-hidden-layer feedforward neural network predicting the policy mean, *µ*_*ϕ*_, and covariance, Σ_*ϕ*_, by taking a cue as input (i.e., the cue encoding different targets in the tasks). In this setting, the actions sampled from the policy were two-dimensional, consisting of the two joint angles controlling the arm position in xy-coordinates. Finally, in the line drawing task, the (Gaussian) policy parametrization consists of a recurrent neural network. This is because the policy has to maintain a memory of the initial cue, which is only presented at the start of the task. We speculate that this (recurrent) policy may be encoded in the motor cortex, which has previously been associated with recurrent neural networks ^127^.

### Motor learning tasks

Motor learning tasks aim to find the policy parameters, *ϕ*, that minimize sensory errors, *e*, while maximizing rewards, *r*. For each trial *t*, the sensory error is computed as the squared distance between the sensory outcome, *y*_*t*_ (e.g., the arm position), and the location of the target, denoted as *y*^∗^:

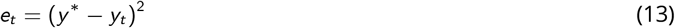

Thus, the minimum error is achieved when the arm reaches the target location, *y*^∗^. Conversely, the reward is modeled in two different ways depending on the motor task:

1. In tasks that provide an explicit binary reward e.g., ^2,19,84^, the reward is computed as a binary success/failure signal, reflecting whether the task is successfully completed. We compute this binary success/failure reward signal as follows:

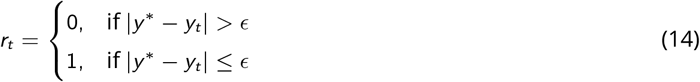

where *ϵ* denotes a minimum accuracy threshold required to consider the task successfully completed.
2. In tasks without an explicit binary success/failure reward signal, we model rewards implicitly to reflect the evaluation of the sensory outcome. In this case, we treat rewards as negative sensory errors, which need to be maximized:

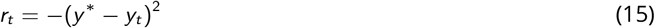

reflecting the fact that achieving smaller sensory errors leads to higher (implicit) rewards.

It is important to stress that our (action-gradient) learning framework does not assume RBL and EBL can only interact when there is a separate explicit reward on top of sensory feedback. Conversely, the same sensory feedback with no explicit reward can trigger both RBL and EBL via their respective action gradient computations.

### Modelling Izawa and Shadmehr arm reaching tasks

We employ an endpoint arm reaching task to reproduce Izawa and Shadmehr ^2^ ‘s results on motor variability and motor generalization under RBL and EBL. Following previous models of reaching adaptation e.g., ^2,27,35^, we model the reaching dynamics based on a linear motor model, where the terminal arm position, *y*, is related to the action, *a* (i.e., reaching angle), as follows:

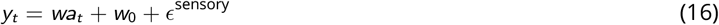

where *{w, w*_0_*}* are randomly initialized, *a*_*t*_ denotes the action sampled from the policy, and 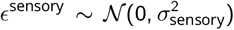 controls the amount of sensory noise over the terminal reaching position. The task follows the exact same setup as Izawa and Shadmehr ^2^, where the model had to reach for a target location, *y*^∗^, under a visuomotor rotation that was gradually increased over 320 reaching trials (i.e., 1 degree increase every 40 trials up to 8 degrees of rotational perturbation) ^2^. To model baseline reaching, we pretrain a policy to reach under normal conditions. Note that this step is not necessary with human participants, since they already know how to reach under normal conditions. Next, we assess the policy’s ability to adapt to the increasing rotational perturbation under RBL, EBL, and a mixture of the two. To assess motor generalization, we also follow the exact same setup as Izawa and Shadmehr ^2^, where the policy generalization performance was tested on novel targets located at [−30, −20, −10, 0, 10, 20, 30] degrees.

### Modelling the visuomotor rotation and reversal tasks

Both tasks involved three phases: 1) an initial baseline phase with no perturbation, 2) a perturbation phase (i.e., where the rotation or reversal perturbation was applied), and 3) a washout phase, where the perturbation is (suddenly) removed. Before the baseline phase, we pretrained a policy to perform the task under normal conditions (i.e., without the perturbation) to establish baseline performance. Note that this step is not necessary with human (adult) participants, since they already know how to perform the task under normal conditions (e.g., they know how to reach with their arm).

For the visuomotor rotation task, we used the same task setup as Tseng et al. ^52^, where we modeled the endpoint arm reaches based on the two-joint kinematic arm model described in the “Two-joint arm model” section. Briefly, the agent had to reach for 3 different target locations placed at angles of [−45°, 0°, 45°] relative to the initial arm position. During the perturbation phase, the visual outcomes in xy-coordinates were rotated by 30° relative to their true position, and the agent had to learn to adapt its reaches to correct for the rotation. The residual error was calculated as the average reaching error (i.e., in terms of the Euclidean distance from the target) over the last 25 trials of the adaptation phase, reflecting the same error measure used by Tseng et al. ^52^ for the experimental data. The interference task included an additional counter-rotation phase, in which an equal-magnitude rotation of opposite sign was applied before washout to disrupt retention of the initially learned rotation and abolish savings.

In the visuomotor reversal tasks, we used the exact same task setup as Gutierrez-Garralda et al. ^62^, where we model dart throwing based on a simple linear model similar to the one described above. The actions denoted the angle direction at which the dart was thrown. For the reversal phase, we introduced a reversal of the perceived angular position of the target by 11.31°, replicating Gutierrez-Garralda et al. ^62^. The adaptation measure was defined as the mean difference between the final throw and the first throw of the perturbation condition, reflecting the same adaptation measure used by Gutierrez-Garralda et al. ^62^ for the experimental data. For the model predictions in Fig. 2d, the residual error was computed as the angular distance from the target angle for the last trial of the adaptation phase (i.e., how much the agent still had to adapt to successfully solve the task). Further experimental details can be found in Gutierrez-Garralda et al. ^62^.

### Targeted endpoint reaching task under dysfunctional cerebellar predictions

For this investigation, we modeled targeted arm reaches based on the two-joint kinematic arm model described in the “Two-joint arm model” section. The state space **s** ∈ ℝ^2^ consisted of the Cartesian coordinates (*x, y*) of the endpoint arm position, while the action space **a** ∈ ℝ^2^ corresponded to the joint angles (i.e., upper arm and forearm). The maximum reach radius was defined by the total arm length (*L*_large_ = *l*_1_ + *l*_2_), and the minimum inner radius was set by the upper arm length (*L*_small_ = *l*_1_) (see the “Two-joint arm model” section for the exact values). The initial hand position (*x*_0_, *y*_0_) was fixed at the midpoint of the reaching workspace:

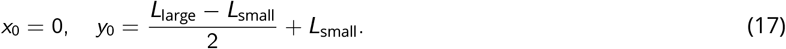

Endpoint targets were distributed uniformly along an 8-target center-out radial layout (*n* = 8) with angular intervals of 45° (*θ* ∈ [0, 2*π*)):

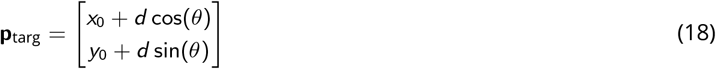

with a fixed target distance *d* = 0.05 m from the origin. Each trial consists of a single-step feedforward reach (*n*_steps_ = 1).

For this task, we did not apply any sensorimotor perturbation to the arm reaches; the policy simply had to reach for the 8 targets under normal conditions. As a first step, we pretrained a policy so that it knows how to reach for the targets (i.e., cerebellar patients usually know how to reach for targets before they experience a cerebellar lesion). Next, we perturbed the cerebellar predictions of the sensitivity derivatives and measured what happened to the reaching endpoints as the policy kept reaching for the targets (i.e., adapting its reaches based on small irreducible errors). This setup aims to replicate cerebellar patients who start performing motor tasks again after their cerebellar lesion. Specifically, the perturbation consisted of switching the sign of one (or more) sensitivity derivatives predicted by the cerebellum. Based on the employed (2D) two-joint kinematic arm model, there were four different sensitivity derivative components that could be perturbed. As mentioned above, the arm takes two-dimensional actions as inputs (i.e., the two joint angles) and outputs a two-dimensional vector encoding the xy-coordinates of the arm endpoint position on a 2-dimensional plane. We report below the four sensitivity derivative components (i.e., the Jacobian matrix of the two-joint arm model):

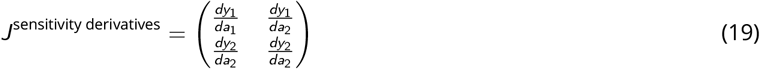

where (*y*_1_, *y*_2_) denote the xy-coordinates of the arm position after taking action (*a*_1_, *a*_2_), encoding the two joint angles (denoted as (*ψ*_1_, *ψ*_2_) in the arm model). Therefore, the cerebellar module had to learn to predict these four components in Eq. 19. In this task, the dysmetria score was computed as the amount of vertical displacement plus the amount of horizontal displacement from each target (i.e., in terms of Euclidean distance), following classical measures of dysmetria see ^61^.

### Discrete reward-based reaching task

For this task, we again modeled arm reaches based on the two-joint arm model described in the “Two-joint arm model” section. Similar to the previous endpoint arm reaching tasks, the goal was to reach for a target, *y*^∗^. However, for this discrete task, no sensory information about the location of *y*^∗^ was available; the task provided only a (binary) success/failure reward signal to guide learning. We compute this binary success/failure reward signal as follows:

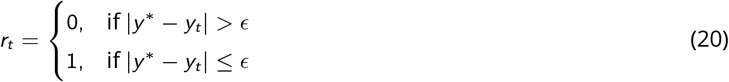

where *ϵ* denotes a minimum accuracy threshold required to consider the task successfully completed.

As shown by previous work e.g., ^2,19,84^, this type of task setup isolates the contribution of (dopamine-driven) RBL to motor learning since (cerebellar-driven) EBL cannot be performed in the absence of sensory feedback, *y* (i.e., one can compute neither the sensory error, *e* = (*y*^∗^ − *y*)^2^, nor the cerebellar-dependent sensitivity derivatives, ^*dy*^, both of which depend on sensory feedback, *y*). Additionally, we model the degree of dopamine deficiency by scaling the reward prediction errors by a factor *γ*:

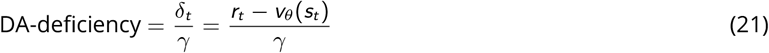

where *γ* ∈ *{*1, 10, 100*}*, representing 3 orders of magnitude of dopamine deficiency, with *γ* = 1 indicating healthy levels of dopamine. Note that this modeling choice is motivated by evidence that phasic dopamine activity encodes reward prediction errors, implying that an overall reduction in (phasic) dopamine activity (e.g., in Parkinson’s disease) should reflect “scaled-down” reward prediction errors ^42,124,125^.

### Drift-diffusion model

To relate action uncertainty to movement onset time, we implemented a simplified single-boundary drift–diffusion model (DDM) in which decision time is governed by stochastic evidence accumulation toward a fixed threshold. In this formulation, the drift rate is modulated by the uncertainty of the learned action policy, operationalized as the standard deviation of the Gaussian policy (*σ*_*a*_). This choice is motivated by empirical and theoretical work linking increased motor and decision uncertainty to delayed action initiation and prolonged reaction times ^80–82^. Specifically, higher uncertainty is assumed to reduce the effective accumulation rate of decision evidence, consistent with normative accounts of confidence-dependent decision dynamics. We define the drift rate *v* as an exponentially decaying function of action uncertainty, *σ*_*a*_ :

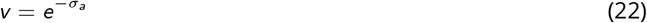

As previously defined, the action uncertainty is determined by the policy-estimated variance 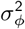 plus a fixed amount of irreducible noise, 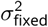 (i.e.,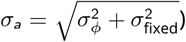). We modeled movement onset time *T* as the time required for the accumulated evidence to reach a fixed decision boundary *κ*:

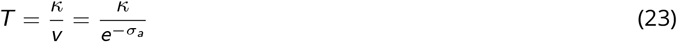

Under this formulation, increased action uncertainty decreases the drift rate, thereby increasing the time required to reach the decision threshold and producing longer movement onset latencies. This is consistent with canonical drift–diffusion formulations of decision time in perceptual and motor decision-making tasks ^80–83^.

### Modelling the savings and interference tasks

To investigate the computational mechanisms underlying savings and interference during motor adaptation, we simulated visuomotor rotation paradigms that alter visual feedback relative to true hand movement using the two-joint kinematic arm model described in the “Two-joint arm model” section. A visuomotor perturbation was imposed by rotating the trajectory of the end-effector by a given angle *θ*_rot_:

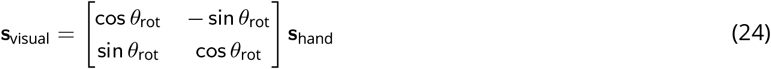

To quantify savings, we evaluated the model’s ability to accelerate relearning following a washout period. The agent first performed baseline trials with visual feedback (*θ*_rot_ = 0°), followed by an initial adaptation block under a counterclockwise perturbation (*θ*_rot_ = +30°). The perturbation was then removed during a washout phase until movement trajectories returned to baseline performance. Crucially, the model was subsequently re-exposed to the same +30° rotation. Savings were demonstrated by comparing the adaptation rate between the initial exposure and the secondary exposure. Accelerated error reduction during early trials of the secondary exposure relative to initial adaptation served as the hallmark of savings.

To evaluate interference, we examined whether learning an opposing perturbation disrupts the retention and subsequent recall of a previously learned motor task. Following initial adaptation to the +30° perturbation (*θ*_rot_ = +30°), the agent was immediately transferred to an opposing clockwise rotation (*θ*_rot_ = −30°), subjected to a washout phase, and then retested on the original rotation. Interference was determined by comparing adaptation rates across the first and final exposures; specifically, complete interference was defined by the network requiring the same number of trials to adapt to the positive rotation during re-test as it did during initial learning, demonstrating that exposure to the opposing rotation completely abolished savings.

### Dopamine-driven offline replay and memory consolidation

To test whether dopamine-dependent reward-based learning (RBL) can support offline memory consolidation, we used a two-phase endpoint reaching paradigm consisting of an initial online learning period followed by a post-learning retention period. To model replay-based consolidation during this retention period, we stored task transitions in a circular memory buffer of fixed capacity *N* during the learning period, implemented as

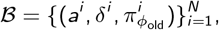

where *a*^*i*^ is the sampled action, δ^*i*^ is the reward-prediction error, and 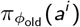 is the action probability under the policy that generated the experience. This is in line with classical theories of hippocampus-driven memory replay in humans ^128,129^. During the post-learning retention period, there is no online learning, and previously stored task transitions are replayed from memory in an offline learning fashion. The off-policy policy gradient update ^97^ provides a principled way to extend the RBL (synaptic) weight update to this offline setting. This extension is achieved by adding an importance-sampling ratio *ρ*_*t*_, defined as

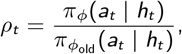

to the standard RBL (synaptic) weight update:

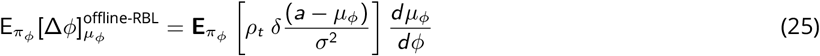

reported here for the policy mean (a similar update can be derived for the policy standard deviation, *σ*_*ϕ*_). Next, to capture forgetting during the post-learning retention period, we additionally introduced a standard multiplicative synaptic decay on the policy parameters encoding the learned actions e.g., ^50^. Specifically, this was implemented as

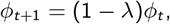

where *λ* is the decay rate. During the post-learning retention period, when offline memory replay was enabled, decay and replay acted together:

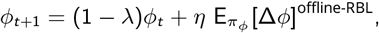

so we could investigate the effect of memory replay in counteracting parameter drift and preserving the learned policy over time. We evaluated retention periodically throughout the post-learning retention phase to quantify the extent to which offline dopamine-driven replay stabilized performance (i.e., via the offline-RBL synaptic weight update above). This analysis was conducted separately for the RBL-acquired and EBL-acquired policies, allowing us to test the prediction that dopamine-dependent replay can also consolidate motor memories acquired via EBL when RBL and EBL affect the same policy parameters ^18,31–34,128^.

### Line drawing task

In the line drawing task, a recurrent (Gaussian) policy has to learn to control the two-joint arm model described in the “Two-joint arm model” section to draw six (target) straight lines, each over 10 time steps, *t*. An initial cue indicates which of the six (target) lines the policy has to draw on any given trial. Crucially, this cue is only provided at the initial time step, *t* = 0. Therefore, the policy has to recall the correct cue over the following 9 steps in order to draw the correct line. This task aims to investigate de novo learning, requiring the policy to learn a complex task completely from scratch rather than through an adaptation paradigm ^48,49^. In this task, the reward is computed as follows:

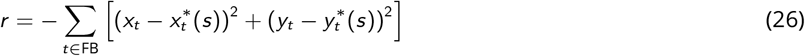

where 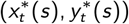 denotes the target location at time *t* to draw the corresponding target straight line given the initial cue, *s*, while FB indicates the set of time steps at which feedback is provided (i.e., controlling the sensory temporal feedback). In order to compute the ‘directed’ sensory error component, 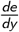, of the EBL action gradient, we assume the brain can directly differentiate the reward (function) relative to *y*. We consider this plausible since, in this task, the reward is again a simple (squared) distance in visual space. Alternatively, a (differentiable) reward function may be learned from practice. As in previous tasks, we assume the cerebellar module learns to predict the sensitivity derivatives, 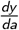, of the EBL action gradient. To study how varying degrees of cerebellar proficiency influence task performance, we defined three levels of cerebellar expertise: *novice* (untrained), *intermediate* (trained to ~50% task accuracy), and *expert* (fully trained). For each condition, the cerebellar module was pretrained accordingly and then frozen, while the remainder of the model was trained from scratch. The optimal *β* was selected via a grid search over [0, 1] (step size 0.2), selecting the value that yielded the highest task accuracy averaged across five random seeds. Cerebellar prediction error was quantified as the mean squared error between the predicted and ground-truth 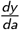 for each sensitivity derivative component, capturing how well the cerebellar module approximates the sensitivity derivatives critical for effective EBL.

### Two-joint arm model

To model arm reaches, we employ a 2-dimensional (non-linear) kinematic arm model ^75^:

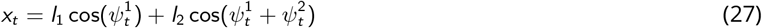

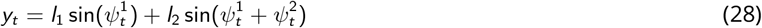

where (*x*_*t*_, *y*_*t*_) denotes the arm position in xy-coordinates at trial *t* (i.e., modelling planar reaching movements). The variables 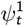 and 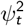 respectively represent the angles of the shoulder and elbow joints at trial *t*, while *l*_1_ and *l*_2_ respectively denote the (constant) upper and lower arm lengths set to standard values *l*_1_ = 0.3348 and *l*_2_ = 0.4572; see ^75^. Therefore, at any given trial *t*, the policy has to provide the two corresponding joint angles to control the position of the arm in space,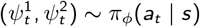.

### Cerebellar module

In the system-level circuits described in the main text, we assume the cerebellum learns to predict the sensitivity derivatives, 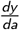, of the EBL action gradient e.g., ^12^. In line with previous work ^48,49^, we represent the cerebellum as a one-hidden-layer feedforward neural network. This cerebellar network is trained to predict the sensitivity derivatives, 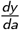, given the current action, *a* (motor command), and the corresponding (sensory) outcome, *y*, at the end of each trial (see Fig. S2 for a visual representation). These predicted sensitivity derivatives, 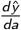, are successively combined with the corresponding ‘directed’ sensory error component, 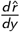 (i.e., coming from sensory cortical areas), to compute the EBL action gradient.

In our experiments, we assume the cerebellar network is provided with the target sensitivity derivatives, 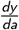. Hence, we train the cerebellar network using a mean squared error loss between the network predictions and the target sensitivity derivatives for each task:

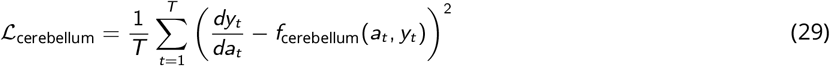

for *T* trials and *f*_cerebellum_(*a*_*t*_, *y*_*t*_) representing the cerebellar network. We followed this procedure for simplicity, as explaining how the cerebellum learns the sensitivity derivatives was beyond the scope of this paper.

Nevertheless, as shown in Fig. S2, we speculate that in the brain, the cerebellum may learn these quantities based on finite-difference methods of gradient estimation see ^130^. Specifically, the cortex could keep track of small changes in (sensory) outcomes, Δ*y*, relative to small changes in motor commands (actions), Δ*a*. The cortex could send this information to the inferior olive, which in turn could compute a finite-difference estimate of the sensitivity derivatives (i.e., by computing the ratio 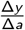), providing this as a teaching signal to the cerebellum in line with well-known mechanisms of the inferior olive providing a teaching signal to the cerebellum; see ^47,131–133^.

Finally, to enable the cerebellar network to model contextual effects in the savings and interference experiments, we modeled context-dependent information by augmenting the cerebellar network with a separate input pathway for each context, as in Pemberton et al. ^50^, where each context is mapped to its own dedicated input pathway. In practice, since our savings/interference experiments involved an unperturbed and a perturbed context, this resulted in two separate input pathways, one per context. This design fully separated the input space such that each context was associated with a non-overlapping set of inputs, preventing overlap between context representations and minimizing interference.

### Action-gradient update implementation

In order to compute the ‘directed’ sensory error component, 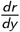, of the EBL action gradient, we directly differentiate the reward (function) in Eq. (15) relative to the observed sensory outcome *y*, to obtain an estimate 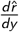. We consider this plausible in the brain since, in the tested tasks, the reward is a simple (squared) distance between the target, *y*^∗^, and the movement outcome, *y*, in visual space (i.e., Eq. 13), suggesting the brain may directly estimate 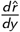 from visual information. Alternatively, the brain may learn a (differentiable) reward function for more complex rewards. Conversely, we assume the sensitivity derivatives, 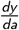, of the EBL action gradient are estimated by a cerebellar module (see “Cerebellar module” section). For instance, in the RL literature, the terms 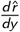 and 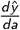 are typically estimated by differentiating through a learned model of the transitions, 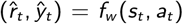, for an initial state *s* i.e., model-based policy gradient; see ^40^. Finally, to correctly estimate the product between the mixed action gradient in Eq. (2) and the term driving synaptic plasticity, 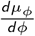, we employ the reparameterization trick. We need this because the policy is Gaussian, implying the relationship between actions, *a*, and the policy parameter, *µ*_*ϕ*_, is stochastic ^134^.

We also report here the ‘mixed’ action-gradient update for the variance parameter of the Gaussian policy,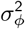 (reported here for the 1-dimensional case for simplicity):

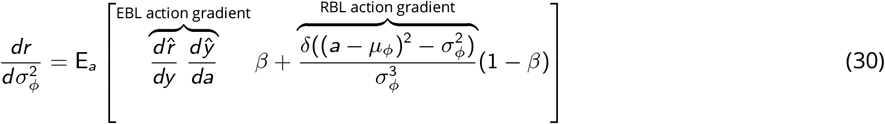

Note that the RBL action gradient on average increases the (policy) variance parameter whenever the reward prediction error is negative (i.e., δ *<* 0), while decreasing it whenever the reward prediction error is positive (i.e., δ *>* 0). Hence, under RBL, we should expect higher motor variability whenever we do not encounter rewards (i.e., leading to negative reward prediction errors) and vice versa.

### Code repository

All the code is available at https://github.com/michele1993/ActionGradients

## Supplementary material

### Implausibility of RBL and EBL training separate policies

In this section, we want to investiagte the possibility that the brain possesses two separate policies for RBL and EBL (e.g., one stored in the cerebellum and another one stored in the basal ganglia), which are only combined at the action selection stage e.g., ^2,16^. In particular, we assess two main ways in which the brain may combine separate RBL and EBL policies. In the first case, the output of the EBL and of the RBL policies could be summed to give the current action (i.e., *a* = *a*^EBL^ + *a*^RBL^). In the second case, the two policy outputs could be combined through a weighted sum, enabling to prioritise one policy over the other when selecting the current action (i.e., *a* = *α a*^EBL^ + (1 − *α*) *a*^RBL^). For instance, in the presence of reward signals, but no sensory feedback, the brain could exclusively rely on the (most updated) RBL policy (i.e., *α* = 0). Conversely, in the presence of optimal sensory feedback, the brain could prioritise the EBL policy (i.e., *α* = 1). Note, this weighted-sum case is in principle similar to what we proposed in eq. 2, but where the weighted sum occurs at the level of actions (based on two separate polices) instead of at level of learning signals (i.e., action gradient). We test these two-policy set-ups in a targeted arm reaching task, where the policy needs to learn to reach for a target location under optimal conditions (i.e., minimal sensory noise and no perturbations). This is the exact same reaching task described in “Izawa and Shadmer arm reaching tasks” Methods section, but without introducing any rotational perturbation. The policy simply has learn to output the correct reaching angles to reach a series of targets across several trials. The targets are placed at different angles from the starting point, thus requiring the policy to learn a different reaching angle for each target (see “Izawa and Shadmer arm reaching tasks” in Methods).

Fig. S3a shows that combining the two policies through a weighted sum (i.e., *α* − weighting) leads to better performance than performing a simple sum of the two polices’ outputs (i.e., No *α* − weighting). This is because for the given task with optimal sensory feedback, EBL seems to be the optimal learning strategy and, the weighted-sum approach (*α* − weighting) is able to suppress the contribution of the (sub-optimal) RBL policy (i.e., by setting *α* = 1). The RBL policy cannot be suppressed in the simple-sum approach (No *α* − weighting), negatively affecting the performance. Furthermore, Fig. S3b shows that combining the two policies through a simple sum leads to higher motor variability than performing the weighted sum, since a simple-sum approach has not mechanism to suppress the extra motor variability induced by the RBL policy when clear sensory information is available. Importantly, this implies that without assuming some weighted combination of RBL and EBL contributions, it is impossible to explain Izawa and Shadmehr ^2^ findings that (human) motor variability changes depending on the quality of the sensory feedback ^2^. Therefore, if the brain holds two separate policies for RBL and EBL, then it likely relies on a weighted sum of the two polices (i.e., *α* − weighting), driving Izawa and Shadmehr ^2^ findings. However, here, we show that a two-policy weighted-sum approach (*α* − weighting) also suffer from a fundamental issue: the two polices fall out of synchrony whenever there is a change in the quality of sensory information affecting the weight. To show this, we assume a reaching task where, at the start, only rewards are provided as feedback with no sensory information (i.e., *α* = 0). After some initial trials, we either suddenly (Fig. S3c) or gradually (Fig. S3d) introduce sensory feedback (i.e., marked by an increase in *α*), thus enabling the EBL policy to also contribute to the actions. Crucially, Fig. S3c-d shows that whenever there is a change in the sensory feedback (i.e., altering the contribution of each policy), the performance suddenly worsens. This occurs because the two policies are no longer synchronized. Conversely, fig. S3c,d show our proposed action gradient framework (green) has no issue in dealing with a sudden or gradual change in the quality of sensory information, supporting learning with no performance worsening. Crucially, the brain is exposed to constant changes to the quality of sensory feedback in the external world. Additionally, performance should likely improve if better sensory feedback becomes available. Therefore, we believe it is implausible the brain holds two separate policies, one trained with EBL and another trained by RBL. We think it is more plausible the brain combines EBL and RBL at the level of teaching signals instead of at the level of actions (i.e., as modelled by our action gradient framework).

### Sensory uncertainty drives the optimal combination of cerebellar and dopamine feedback

A key prediction of our action gradient framework is that the brain should rely on a (*β*-) weighted combination of the cerebellar-driven (EBL) and the dopamine-driven (RBL) learning signals. The degree of combination should partially depend on the amount of uncertainly over the sensory information, *y* (i.e., the higher the uncertainty, the lower reliance on cerebellar-dependent EBL). This is because when the sensory outcome *y* cannot be estimated correctly, the cerbellar driven EBL action gradient provides a noisy teaching signal to drive learning. To probe this prediction, we investigate which RBL-EBL action gradient combinations lead to optimal learning performance across different levels of noise over the sensory information, paired with a “noise-free” success/failure reward signals (i.e., in terms of the optimal *β* combination of the RBL and EBL action gradients in eq. 2). For instance, this setting could be equivalent to a tennis game where you can estimate the tennis bal position after a shot with different degrees of accuracy (i.e., different levels of sensory noise), but you are told if you scored a point (i.e., “noise-free” binary reward signal). To model this more complex setting, we employ a neural network-based simulation of motor dynamics in a 3-dimensional outcome space controlled via a 10 dimensional action space (e.g., a small population of neurons encoding motor commands). The goal of the task is the same as previous motor tasks, learning a policy, *π*_*ϕ*_, that achieve a desired outcome, *y*^∗^. The exact experimental details are reported in the next paragraph. Fig. S4 shows the optimal action gradient should reflect a combination of both the cerebellar-driven (EBL) and the dopamine-driven (RBL) learning signals (action gradients) across most levels of sensory noises (i.e., 0 *<* Optimal *β <* 1 for most sensory noise levels). This indicates it is beneficial to combine EBL and RBL learning signals together rather than relying on one or the other. Crucially, as the sensory noise increases, the contribution of the cerebellar-driven EBL learning signals should rapidly decrease in favor of higher contributions from the dopamine-driven RBL learning signals. This is because as the sensory noise increases, the accuracy of the cerebellar-driven EBL learning signals decreases, since they depend on sensory errors. However, this inaccuracy can partially be mitigated by a greater contribution of the dopamine-driven RBL learning signals, providing a mechanism to learn under high sensory noise. In the ‘real’ world sensory information is encountered with different degrees of noise, implying the brain may greatly benefit from combining the reward- and error-based learning signals, reflecting close basal ganglia and cerebellar interactions e.g., ^21,24–26^. This prediction should experimentally be validated in a motor learning task where the quality of the sensory feedback is varied continuously together with reward signals. As the the quality of the sensory outcome decreases learning should move from cerebellar-driven towards dopamine-driven. Based on what we showed in the motor variability experiment, this partial switch from cerebellar-to dopamine-driven learning should be observable behaviorally by measuring any increase in motor variability, which would indicate learning is increasingly driven by dopamine. This provides a simple avenue for experimentally validating an important prediction of the proposed learning framework.

This task aimed to reproduce the complex non-linear and high-dimensional relation between actions, *a*, and sensory outcomes, *y*, during arm reaches in a 3-dimensional space. To do so, we model the task dynamics with a high-dimensional non-linear feedforward neural network with fixed weights (i.e., fixed task dynamics). This network comprised a single hidden layer of 256 hidden units, taking 10-dimensional actions as input, *a*, and outputting a 3 dimensional outcome, *y* (e.g., xyz-coordiante in space). The 10-dimensional action space can be thought of as encoded by the firing frequency of 10 small population of neurons outputting efferent motor commands. The goal of the task was the same as the other motor tasks, learning a policy, *π*_*ϕ*_, that achieve the desired outcome, *y*^∗^. Specifically, the task comprised of 10 targets, *y*^∗^, which the policy had to achieve. The policy consisted of a single hidden layer feedforward neural network (56 hidden units), taking a cue as input and outputing a Gaussian distribution over the actions (i.e., a Gaussian polciy). The cue served to distinguish the 10 different targets. Similarly to all the other tasks, the reward function, *r*, was the (negative) of the squared euclidean distance between the current output and the target output, (i.e., *r* = −(*y*^∗^ − *y*)^2^). The sensory noise levels were modelled by adding zero mean Gaussian noise of increasing standard deviations to the outcomes, *y*. The optimal *β* was determined by finding the *β* value that lead to learning the policy achieving the highest task reward across an entire training run with a fixed level of sensory noise (i.e., for 5000 trials/episodes).

### Long-Term Effects of Savings and Interference

Beyond the results presented in the main text, we examined whether extending the washout phase would influence the observed savings and interference. In empirical studies, savings are often measured over extended washout periods, such as across different days. This raises the question of whether our model can retain savings over longer durations. To investigate this, we tested whether a prolonged washout period might diminish or obscure these effects. As shown in Figure S5, the patterns of long-term savings and interference remain consistent, demonstrating that our findings are robust to variations in washout duration.

### Impact of Context-Dependent Parallel Fiber Activation on Savings

Building on the findings of ^50^, which align with observed context-dependent activations and plasticity rules in the cerebellum, we focused on cerebellar Parallel Fibers (PFs) that exhibit task-specific activity. The degree of task specificity in PFs is modeled using the PF task overlap: maximal overlap (no context) implies the same set of PFs is engaged across all task contexts, whereas zero overlap (context) indicates entirely distinct PFs for each task. Pemberton et al. ^50^ demonstrated that PF task overlap mediates a tradeoff between learning speed for new tasks and the ability to rapidly revert to previously learned tasks. Specifically, high overlap accelerates initial learning, while distinct PF sets facilitate quick switching between tasks.

In this study, we investigated how PF task overlap influences savings and how these effects change with varying washout periods. We quantified savings by calculating the area between the adaptation and re-adaptation curves (i.e., the second exposure to the same rotation). As shown in Figure S6, context-dependent PF activation enhances savings, particularly with longer washout periods, whereas without contextual differentiation, savings diminish as the washout duration increases.

These results are consistent with expectations. When PFs are context-dependent (0% overlap), distinct fiber sets encode each task, minimizing interference and preserving memory traces during washout, thereby enhancing savings. In contrast, with high PF overlap, the same fibers encode multiple tasks, leading to interference and partial overwriting of prior learning, which diminishes savings.

### Dopamine-driven Learning Alone Does Not Produce Savings

A central claim of this paper is that the computational mechanism underlying savings depends critically on a cerebellar caching process. To isolate the contribution of the dopamine RBL teaching signal, we tested the model behavior when cerebellar involvement was removed. Concretely, this was implemented by setting the *β* parameter to 0, thereby restricting learning entirely to the dopamine-driven pathway.

Under this manipulation, the model retained the capacity to gradually adapt to the perturbation during the initial learning phase. However, critically, it failed to exhibit savings during relearning. As shown in Figure S7, performance during the second exposure to the same perturbation closely mirrors the original acquisition curve. There is no observable acceleration in relearning, indicating the absence of a stored or cached memory trace that could be rapidly re-engaged.

This result demonstrates that RBL, when working alone, is insufficient to account for savings. Although the striatum is capable of supporting gradual, reinforcement-driven adaptation through incremental updating of action values, it lacks a mechanism for rapid memory caching. As a result, when the perturbation is reintroduced, learning must proceed *de novo* rather than being accelerated by access to a previously consolidated internal representation, thereby preventing the emergence of savings. In contrast, when the cerebellar system is active (e.g., *β* = 0.5), the model shows clear evidence of savings, consistent with the hypothesis that the cerebellum provides a memory buffer or internal model that can be rapidly reactivated. Therefore, these findings support the interpretation that savings is not simply a byproduct of repeated reinforcement learning, but instead emerges from a distinct cerebellar-dependent caching mechanism.

---

**Figure S1.**
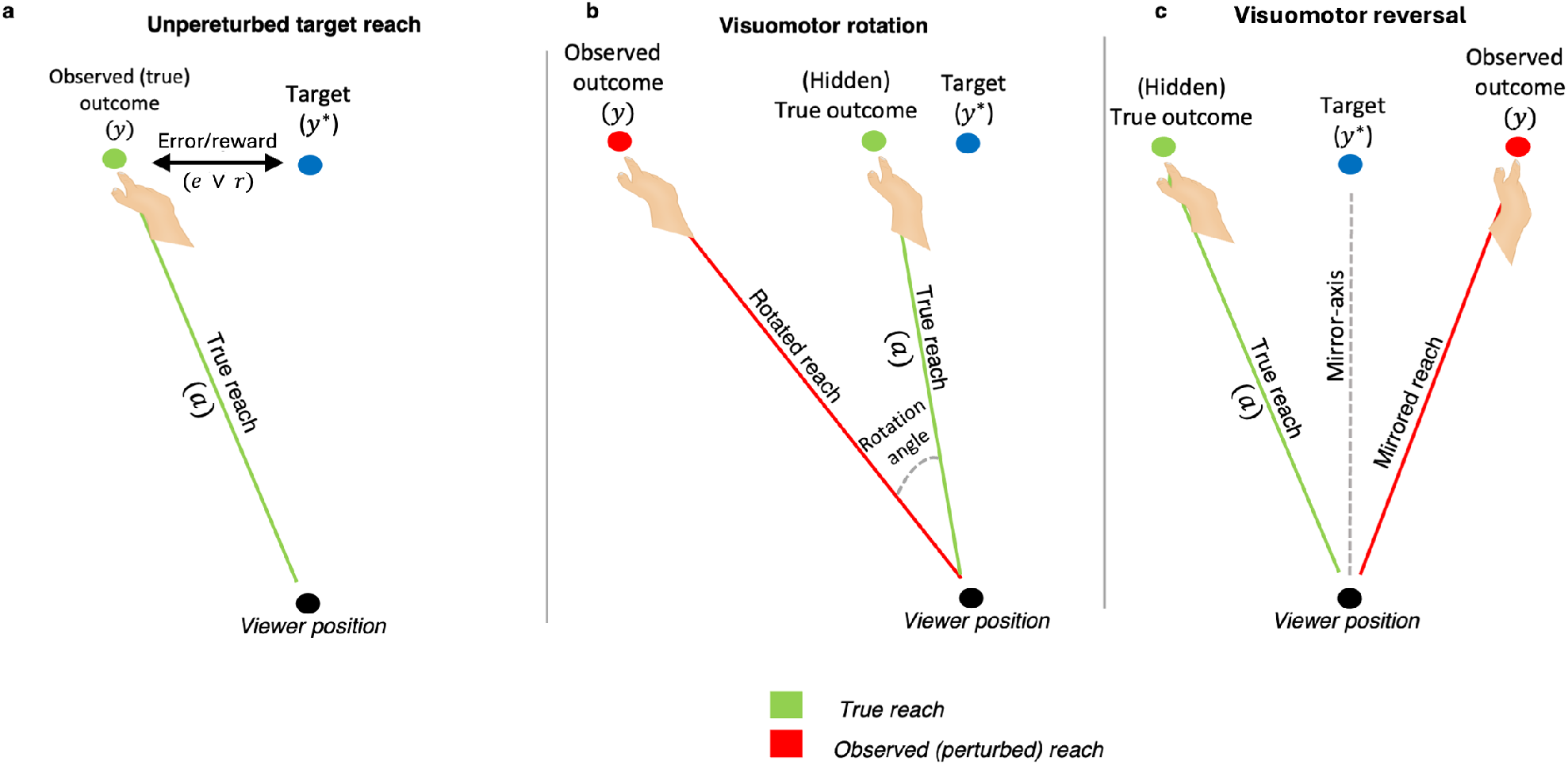
Visuomotor task representations. **a**, Representation of a standard (unperturbed) end-point reaching task where subjects need to reach with their arm for target, *y*^∗^, based on the distance from the observed reaching outcome *y* and the target *y*^∗^, a sensory error and/or implicit reward signal can be estimated. **b**, Representation of a visuomotor rotation end-point reaching task, where the true reaching outcome is hidden and only the rotated outcome is observable. **c**, Representation of a visuomotor reversal end-point reaching task, where the true reaching outcome is hidden and only the mirror reversed outcome is observable. The mirror axis represents the target axis over which the reaching outcomes are mirrored.

**Figure S2.**
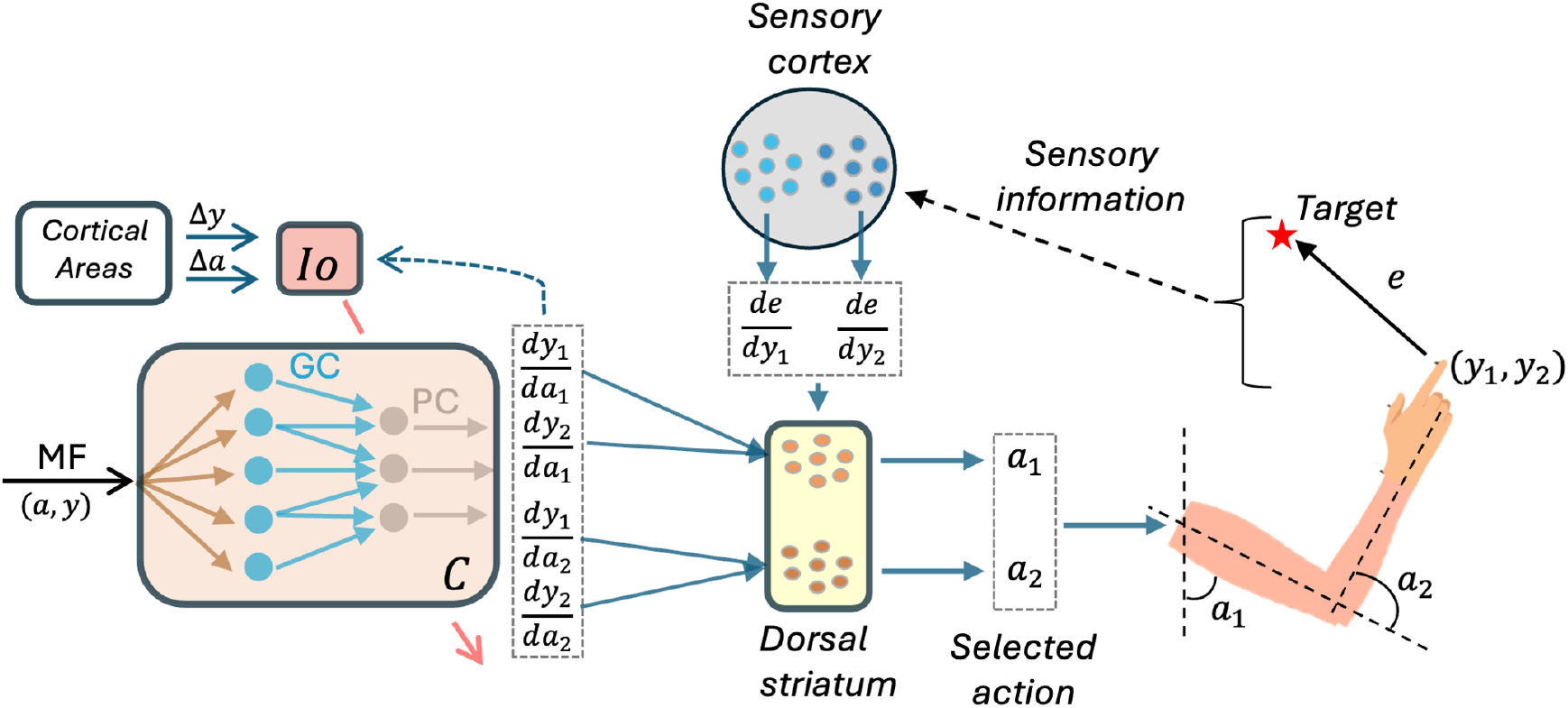
In-depth diagram of the EBL action gradient system-level circuit for an endpoint reaching task on a 2-dimensional place. MF: mossy fibers, C: cerebellum, Io: inferior olive. Representation of the cerebellar network as a 1 hidden-layer feedforward neural network ^48,49^. The mossy fibers (MF) convey the cerebellar input, encoding the motor commands, *a* and the corresponding (sensory) outcomes, *y*, at the end of each trial/step, *t*. Granule cells (GC) represent the hidden layer computations. Finally, Purkinje cells (PC) represent the output layer, providing the sensitivity derivative predictions, 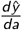. This cerebellar network learns based on the inferior olive (Io) feedback, conveying a mismatch between the (current) cerebellar output and observed changes in (sensory) outcomes (i.e., Δ*y*) relative to the changes in motor commands (i.e., Δ*a*) e.g., exploiting a finite difference gradient estimate ^130^. Here, we assume two different populations of dorsal striatal cells respectively encode the joint angle for the shoulder, *a*_1_, and for the elbow, *a*_2_, controlling the arm location. On a 2-dimensional plane, the arm location is determined by two coordinates, (*y*_1_, *y*_2_), based on which a sensory error, *e*, can be computed, encoding the distance between the desired target and the current arm location. From this sensory error *e*, we assume two different populations of sensory cortical neurons compute the two corresponding ‘directed’ sensory errors, 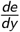, one for each coordinate. These two ‘directed’ sensory errors are then sent back to the dorsal striatum, instructing it how to adjust the selected action, thanks to the cerebellar projections, encoding.

**Figure S3.**
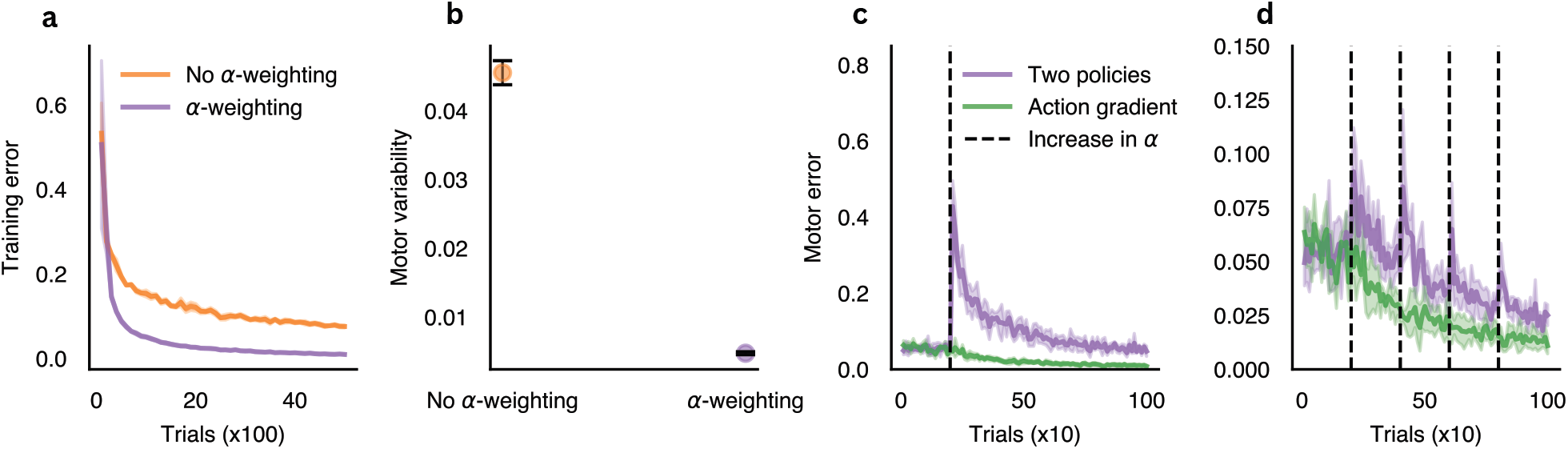
Learning performance for separate EBL and RBL policies. The two policies are combined by either a simple sum (No *α*−weighting) or based on a weighted sum (*α*−weighting). **a**, the weighted-sum approach achieves better learning performance than the simple sum approach. **b**, The weighted-sum approach achieves lower motor variability than the simple sum approach. **c-d**, the weighted-sum approach (purple) suffers from a drop in performance whenever there is a sudden (**c**) or gradual (**d**) change in sensory vs reward feedback, unlike our proposed action gradient approach (green).

**Figure S4.**
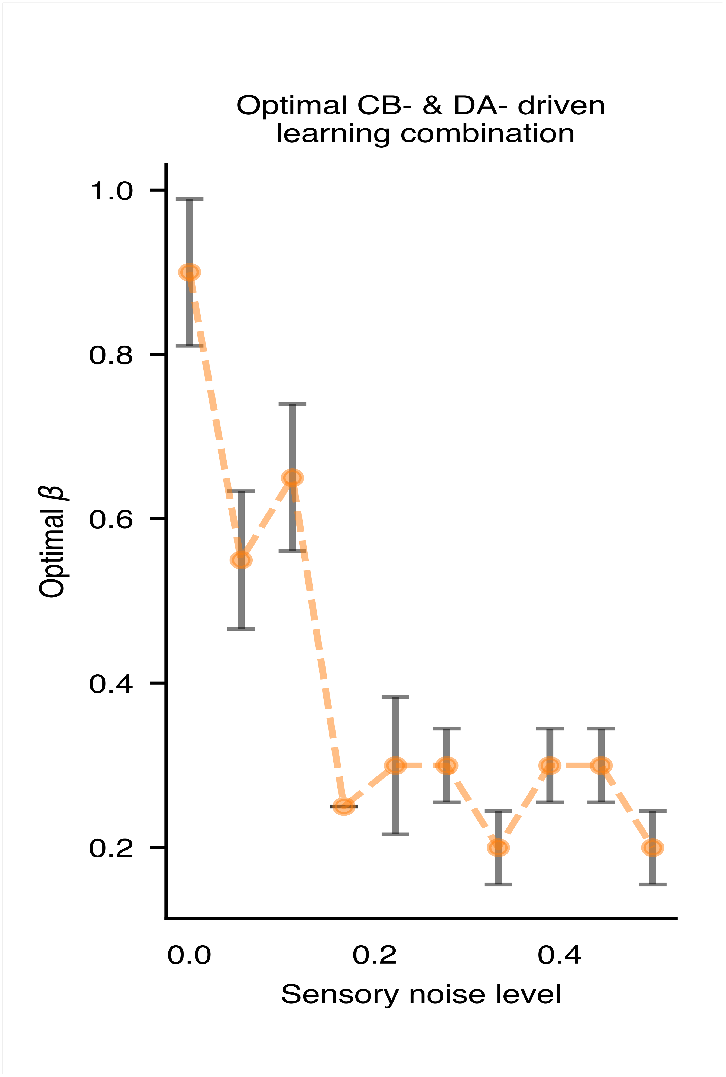
CB: cerebellum, DA: dopamine. Uncertainty over the sensory feedback *y* affects the optimal *β*− combination of RBL and EBL teaching signals (i.e., action gradient)

**Figure S5.**
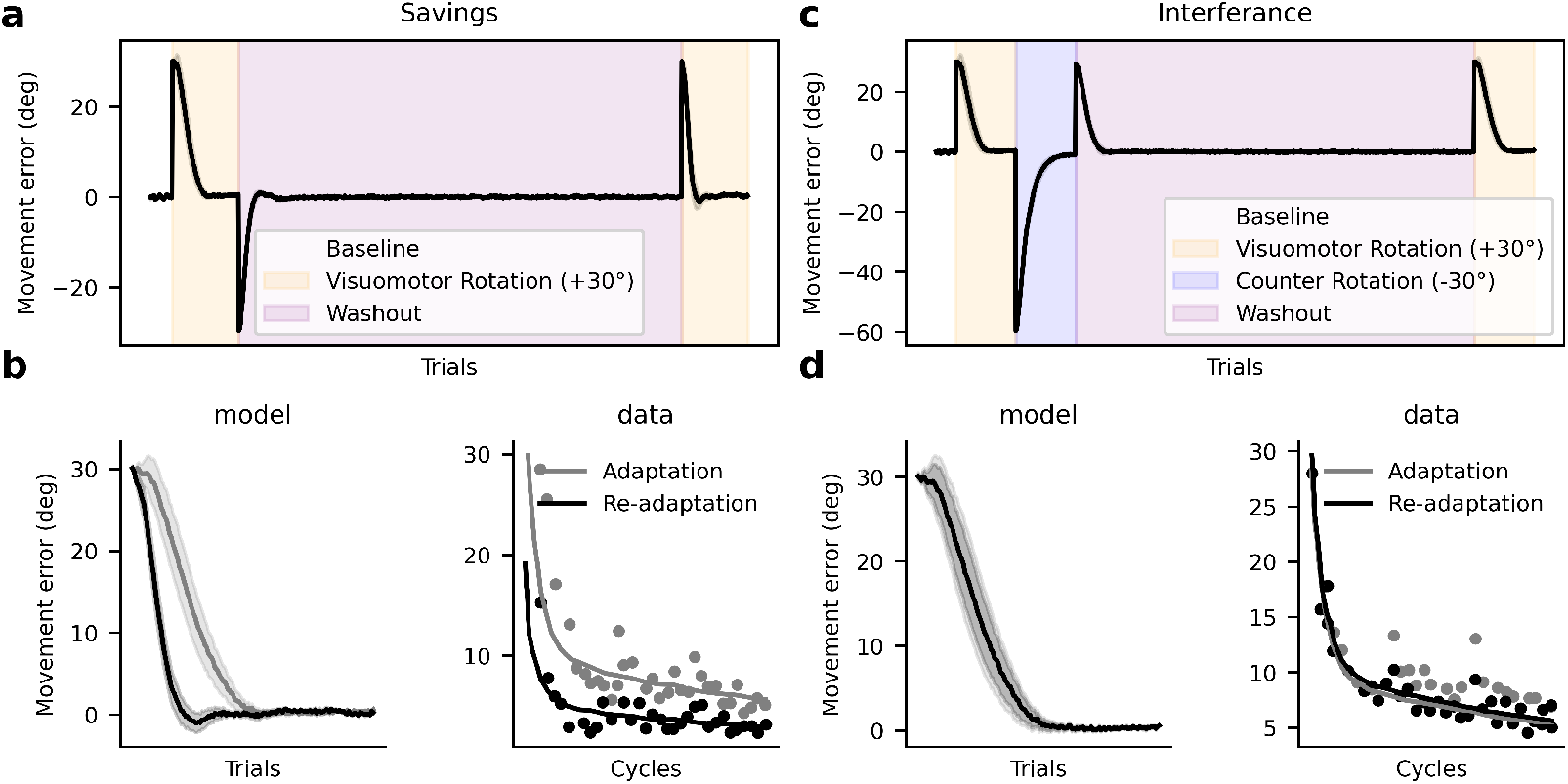
Long-term effects of savings and interference after an extended washout phase. The results demonstrate that both savings and interference are preserved, showing the robustness of these effects over time.

**Figure S6.**
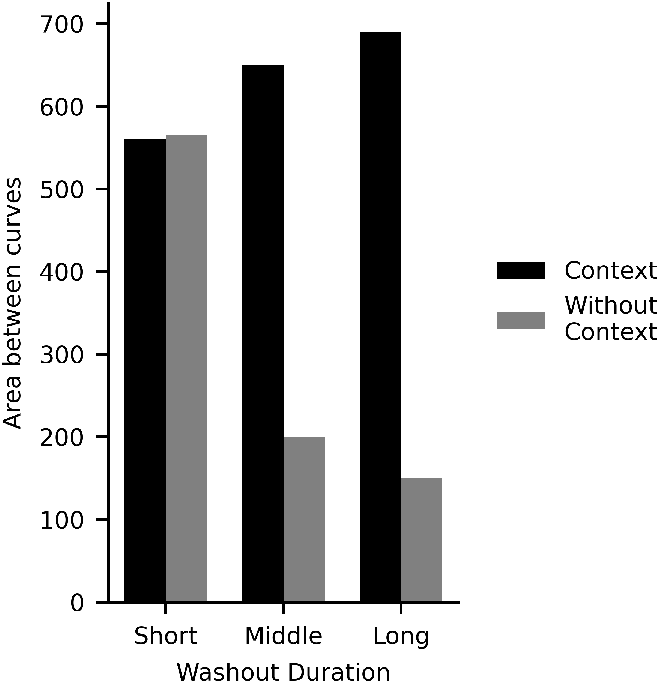
Effect of PF task overlap on savings across washout periods. Context-dependent PF activation (0% overlap) increases savings with longer washout, whereas high overlap (no context) leads to reduced savings. The area between adaptation and re-adaptation curves quantifies the magnitude of savings.

**Figure S7.**
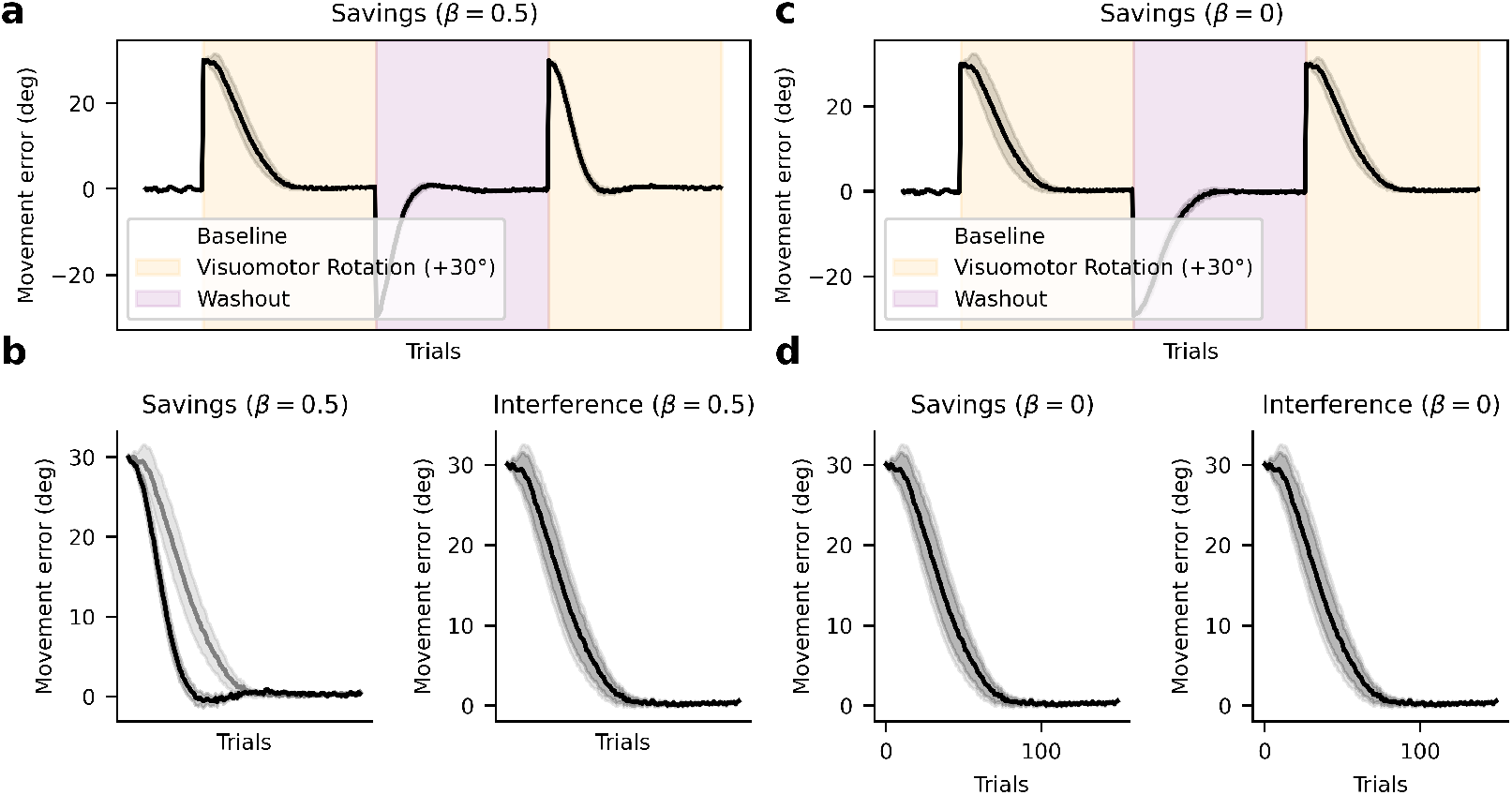
Learning curves when the cerebellar contribution is removed (*β* = 0). Although the model adapts during initial exposure to the perturbation, relearning proceeds at a similar rate, showing no acceleration relative to first learning. The absence of improved performance during the second exposure indicates that striatum-driven learning alone does not produce savings.

**Figure S8.**
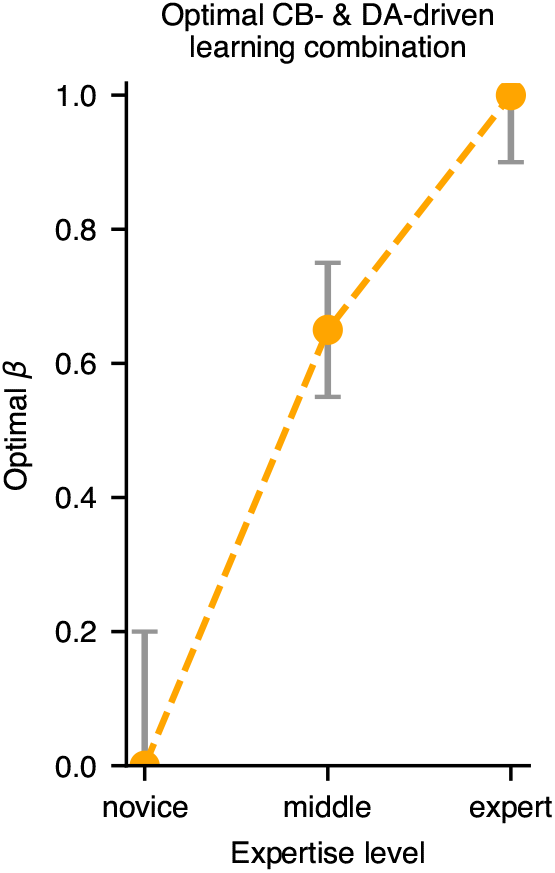
Optimal combination of CB and DA-dependent learning depends on task expertise for a simple Center-Out Reaching Task.

